# Dynamic RNA Polymerase compartments organize the transcription of gene clusters

**DOI:** 10.64898/2026.01.07.698080

**Authors:** Yi-Hui Wang, Hannah L. Hertz, Wen Tang

**Affiliations:** Department of Biological Chemistry and Pharmacology, The Ohio State University, Columbus, OH 43210, USA; Center for RNA Biology, The Ohio State University, Columbus, OH 43210, USA; Ohio State Biochemistry Program, The Ohio State University, Columbus, OH 43210, USA

## Abstract

Spatial organization of transcription machinery is a key regulator of gene expression, yet how different RNA polymerases are organized at a single gene cluster is not well studied. Here, we show that RNA polymerases II and polymerase III form nuclear foci at the 5S ribosomal DNA (rDNA)–Spliced leader 1 (SL1) cluster in *C. elegans*. Within this cluster, polymerase II binds to the SL1 gene, while polymerase III associates with 5S rDNA. Both polymerase foci display dynamic but distinct behaviors within the nucleus. The assembly of these polymerase foci is regulated across the cell cycle. ATTF-6, an AT-hook transcription factor, is essential for polymerases II foci formation but dispensable for polymerases III foci. While Pol III foci are largely resistant to temperature changes, Pol II foci are temperature-sensitive, and their dissolution correlates with reduced SL1 expression. Taken together, these results reveal a spatial and temporal regulation of two RNA polymerases that organize gene cluster transcription.

## Introduction

In eukaryotes, transcription is carried out by three major RNA polymerases, each dedicated to distinct classes of genes. RNA polymerase I (Pol I) transcribes ribosomal RNAs (rRNAs), polymerase II (Pol II) produces messenger RNAs (mRNAs) and many long non-coding RNAs, and RNA polymerase III (Pol III) synthesizes transfer RNAs (tRNAs), 5S rRNA, and other small structural RNAs (Cramer 2019; Roeder 2019; Girbig et al. 2022). Beyond diffuse nucleoplasmic distributions, accumulating evidence indicates that transcription is spatially organized into focal compartments or condensates that coordinate efficiency and specificity (Rippe 2022; Rippe and Papantonis 2025).

RNA Pol I is highly enriched in the nucleolus, a membraneless compartment specialized for rRNA biogenesis (Grummt 2003). In contrast, Pol II is expressed throughout the nucleoplasm. Pol II transcription can occur within discrete “transcription factories” or nuclear foci (Jackson et al. 1993; Wansink et al. 1993). These foci represent sites where multiple genes, transcription factors, and nascent transcripts are locally concentrated, providing a framework for coordinated transcription and RNA processing. Several mechanisms have been proposed to explain their formation. Multivalent interactions among Mediator components, transcription factors, and the intrinsically disordered C-terminal domain of Pol II may promote liquid-liquid phase separation or related microphase-separation processes (Hnisz et al. 2017; Boehning et al. 2018; Cho et al. 2018; Palacio and Taatjes 2022). Alternatively, transcription factors and polymerases may become locally enriched through chromatin-scaffolded recruitment to clustered binding sites, such as those provided by repetitive DNA elements (Erdel and Rippe 2018; Muzzopappa et al. 2022). Chromatin scaffolding and phase separation are not mutually exclusive, and multivalent interactions among chromatin, transcription factors, and polymerases can contribute together to the formation and stabilization of transcriptional foci (McSwiggen et al. 2019; Stortz et al. 2024; Rippe and Papantonis 2025).

Compared to Pol I and Pol II, the spatial organization of Pol III is less understood. Pol III transcribes non-coding RNAs such as tRNAs and 5S rRNAs, both of which are essential for translation. Studies suggest that its activity may concentrate at specific nuclear foci, analogous to Pol II (Pombo et al. 1999). Yet, the spatial relationship between Pol III, Pol II and other transcriptional machineries, and how these polymerase-specific compartments are regulated, remain largely unknown.

Many organisms organize functionally related genes into clusters, including piRNA, histone, and rRNA genomic loci (Blumenthal 1998; Ruby et al. 2006; Duronio and Marzluff 2017; Pastore et al. 2022). In *C. elegans*, the 5S rDNA–spliced leader 1 (SL1) cluster offers a unique opportunity to study how Pol II and Pol III transcriptional programs coexist in close proximity. This cluster consists of more than 100 tandem ∼1 kb repeats on chromosome V, each unit harboring one 5S rDNA gene and one SL1 gene (Ellis et al. 1986; Ding et al. 2022). The 5S rRNA is transcribed by Pol III whose largest subunit is RPC-1 in *C. elegans*, while SL1 is transcribed by Pol II with AMA-1 as its catalytic subunit (Rogalski and Riddle 1988; Bird and Riddle 1989). SL1 encodes a specialized small nuclear RNA (snRNA) required for trans-splicing, a process that appends SL1 to the 5′ ends of more than half of all pre-mRNAs to promote efficient translation initiation (Blumenthal 2012; Yang et al. 2017). The interspersed arrangement of Pol II- and Pol III-transcribed genes provides a repetitive chromatin scaffold that may locally concentrate both transcriptional machineries through chromatin-scaffolded assembly, liquid–liquid phase separation, or a combination of these mechanisms. However, how Pol II and Pol III are organized and differentially regulated within this shared locus remains unknown.

We previously identified ATTF 6 (an AT hook family protein) as a key regulator of several gene clusters in *C. elegans*, including histone, piRNA, and 5S rDNA–SL1 clusters (Wang et al. 2025). ATTF 6 forms discrete nuclear foci at both piRNA and 5S rDNA–SL1 clusters in the germline. Loss of ATTF 6 results in reduced expression of histone mRNAs, SL1 transcripts, and piRNAs (Wang et al. 2025). These findings suggest that ATTF 6 acts broadly to promote transcription at clustered genomic loci, but whether it contributes to the spatial organization of RNA polymerases has not been addressed.

Here, we exploit this dual-polymerase 5S rDNA–SL1 cluster to investigate how two RNA polymerases form spatially organized transcription compartments in vivo. We show that Pol II and Pol III form separate, dynamic nuclear foci at the 5S rDNA–SL1 cluster, each regulated in a cell cycle–dependent manner. Pol II foci require the transcription factor ATTF-6, are sensitive to temperature, and regulate SL1 expression, whereas Pol III foci are less affected by these conditions. Together, our findings reveal that *C. elegans* RNA polymerases assemble into condensate-like compartments at shared gene clusters, uncovering an unexpected layer of spatial organization that coordinates transcription across polymerase systems.

## Results

### RNA Pol II and Pol III form nuclear foci at the 5S rDNA–SL1 cluster

Our recent work showed that ATTF-6, an AT-hook transcription factor, associates with genomic clusters including the piRNA cluster and the 5S rDNA–SL1 cluster in the *C. elegans* gonad (Wang et al. 2025). The gonad is a syncytial tissue in which germ cells proliferate mitotically at the distal end and then enter meiotic pachytene to produce oocytes, which are fertilized to form embryos (Figure S1A) (Pazdernik and Schedl 2013). ATTF-6 forms a large nuclear focus at the piRNA cluster specifically in the early– mid pachytene region, where homologous chromosomes undergo synapsis (Figures S1B-S1G) (Batista et al. 2008; Das et al. 2008; Wang et al. 2025). Furthermore, ATTF-6 forms a smaller and discrete focus at the 5S rDNA–SL1 cluster throughout pachytene and oogenesis (Figures S1B-S1G) (Wang et al. 2025).

Because early embryos undergo rapid cell divisions and execute transcriptional programs different from those of the germline, we asked whether ATTF-6 remains associated with the 5S rDNA–SL1 cluster in embryonic cells. To this end, we performed 5S rDNA FISH (Fluorescence in situ hybridization) followed by immunofluorescence against ATTF-6::3xFLAG using an anti-FLAG antibody (Figures 1A and 1B). ATTF-6::3xFLAG formed discrete nuclear foci in embryos that colocalized with the 5S rDNA– SL1 cluster indicated by 5S rDNA FISH signals (Figures 1A and 1B). These results indicate that ATTF-6 associates with the cluster not only in germ cells but also in embryonic nuclei.

**Figure 1.**
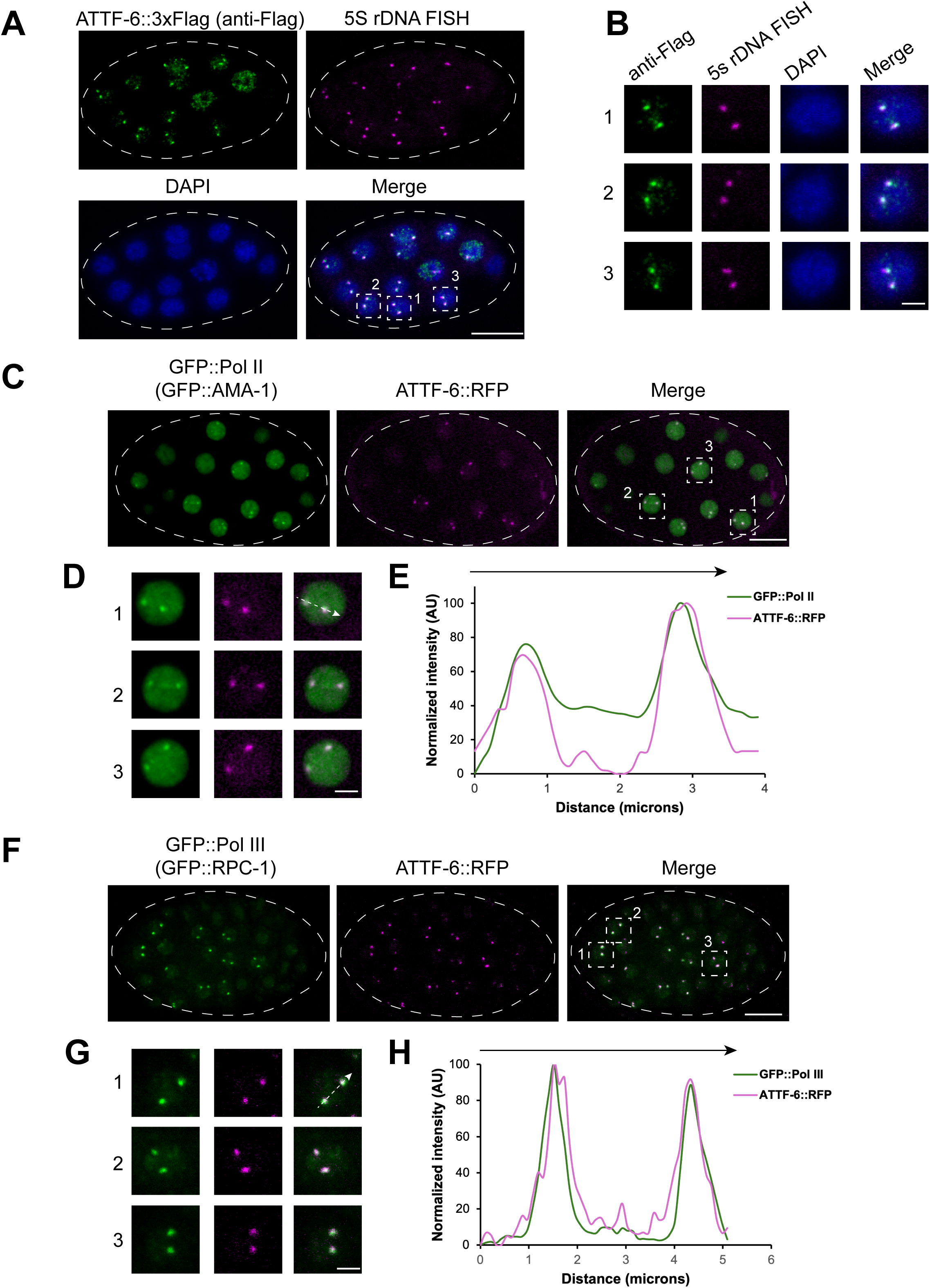
RNA Pol II and Pol III form nuclear foci at the 5S rDNA-SL1 gene cluster. **(A)** Co-localization of 5S rDNA and ATTF-6 in embryo nuclei detected by DNA-FISH and immunostaining. DNA-FISH was performed using Cy5-labeled probes targeting the 5S rDNA gene cluster, and immunostaining was conducted with an anti-FLAG antibody to detect ATTF-6::3×FLAG. Nuclei were visualized with DAPI. Dashed lines outline the embryo. Confocal images (60× objective) are shown as maximum-intensity projections spanning the top layer of nuclei in the fixed embryo. Scale bar: 10 μm. **(B)** Enlarged view of three nuclei within the embryo shown in (**A**). Scale bar: 2 μm. **(C)** GFP::Pol II (GFP:: AMA-1) and ATTF-6::RFP foci in a live embryo. Dashed lines outline the embryo. Confocal images (60× objective) are shown as maximum-intensity projections spanning the top layer of nuclei in the embryo. Scale bar: 10 μm. **(D)** Enlarged view of three nuclei within the embryo shown in (**C**). Scale bar: 2 μm. **(E)** Intensity profile of GFP::Pol II (GFP::AMA-1) and ATTF-6::RFP signals along the dotted arrow in nucleus No1. in (**D**). AU, arbitrary unit. **(F)** GFP::Pol III (GFP::RPC-1) and ATTF-6::RFP foci in a live embryo. Dashed lines outline the embryo. Confocal images (60× objective) are shown as maximum-intensity projections spanning the top layer of nuclei in the embryo. Scale bar: 10 μm. **(G)** Enlarged view of three nuclei within the embryo shown in (**F**). Scale bar: 2 μm. **(H)** Intensity profile of GFP::Pol III (GFP::RPC-1) and ATTF-6::RFP signals along the dotted arrow in nucleus No. 1 in (**G**). AU, arbitrary units.

The 5S rDNA–SL1 cluster is unique in that it contains over 100 tandem copies of the 5S rRNA gene and SL1 snRNA gene (Ellis et al. 1986; Ding et al. 2022). The 5S rRNA, transcribed by Pol III, is an essential component of ribosomes, while SL1, transcribed by Pol II, provides the spliced leader sequence for trans-splicing (Blumenthal 2012; Yang et al. 2017). The presence of two polymerase-dependent transcription units within a highly repetitive genomic region raises the question of whether Pol II and Pol III are locally enriched at this cluster.

To address this, we examined the subcellular localization of GFP::AMA-1 (the largest catalytic subunit of Pol II) and GFP::RPC-1 (the largest catalytic subunit of Pol III) expressed from their endogenous genomic loci. GFP::AMA-1 (hereafter referred to as GFP::Pol II) was broadly distributed throughout the nucleus but showed noticeable enrichment at the small ATTF-6::RFP focus corresponding to the 5S rDNA–SL1 cluster in the gonad (Figure S1B). Line-scan analysis revealed substantial overlap between the fluorescence intensity profiles of ATTF-6::RFP and GFP::Pol II across the small ATTF-6 focus (Figure S1C). Similarly, GFP::RPC-1 (hereafter referred to as GFP::Pol III) was enriched at and overlapped with the small ATTF-6::RFP focus (Figures S1D and S1E). Neither polymerase was enriched at piRNA clusters marked by mCherry::PRDE-1, a piRNA specific transcription factor (Figures S1F and S1G) (Weick et al. 2014; Weng et al. 2019; Wang et al. 2025).

In embryos, confocal microscopy revealed that GFP::Pol II formed several nuclear foci, including prominent two that colocalized with ATTF-6::RFP (Figures 1C and 1D), as confirmed by line-scan intensity profiles (Figure 1E). Similarly, GFP::Pol III formed two prominent foci in embryos that overlapped with ATTF-6 (Figures 1F-1H). These data demonstrate that Pol II and Pol III assemble into nuclear foci at the 5S rDNA–SL1 cluster in both the germline and early embryos.

### Pol II associates with SL1 gene while Pol III associates with 5S rDNA

To further validate the enrichment of polymerases at the 5S rDNA–SL1 cluster, we reanalyzed published ChIP-seq (Chromatin Immunoprecipitation followed by high-throughput sequencing) datasets (Ikegami and Lieb 2013; Wang et al. 2025). In these experiments, ChIP was performed from mixed-stage embryos using antibodies against AMA-1 (Pol II) or RPC-1 (Pol III) (Ikegami and Lieb 2013). ATTF-6 ChIP-seq data were previously generated using anti-Flag antibody in synchronized adults expressing endogenously tagged ATTF-6::3xFlag (Wang et al. 2025). Here we examined ATTF-6, Pol II and Pol III ChIP signals across individual chromosomes including five autosomes and one sex chromosome (chromosome X). Consistent with our previous findings (Wang et al. 2025), genome-wide profiles showed that ATTF-6 binds strongly to the piRNA cluster on Chromosome IV and to the 5S rDNA–SL1 cluster on Chromosome V (Figures 2A and S2). Pol II signals were broadly distributed across the genome, with strong enrichment at several loci including histone gene clusters on chromosomes II and IV. (Figure S2). Notably, the single strongest Pol II and Pol III ChIP peaks across the genome occurred at the 5S rDNA–SL1 locus (Figures 2A, 2B and S2).

**Figure 2.**
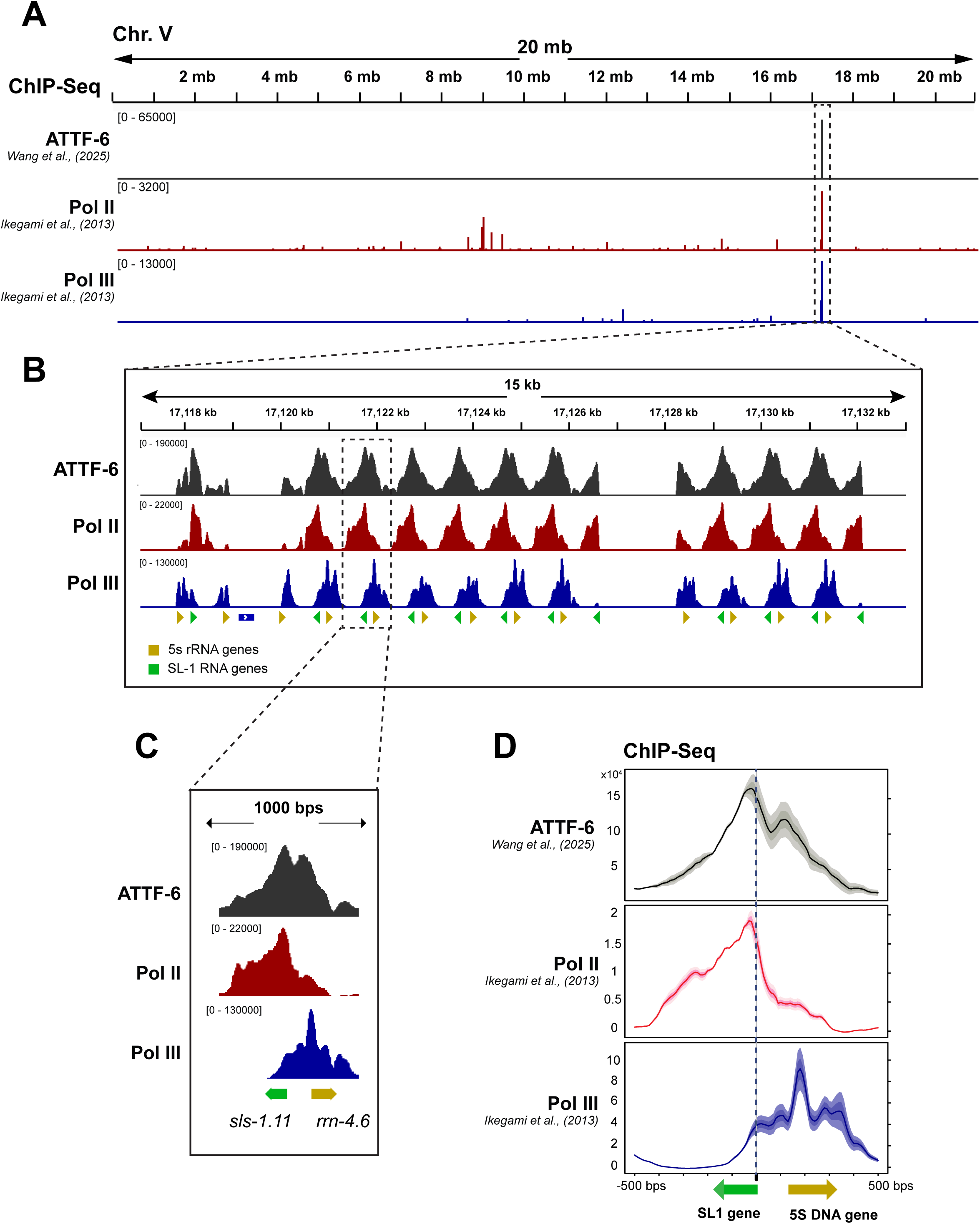
RNA Pol II and Pol III associate with SL1 and 5S rDNA, respectively. **(A)** Browser view of ATTF-6, Pol II (AMA-1), and Pol III (RPC-1) ChIP-seq signals on chromosome V. ATTF-6 ChIP-seq signals were normalized to the control IP (Wang et al. 2025), and AMA-1 and RPC-1 ChIP-signals were normalized to the input (Ikegami and Lieb 2013). The signals represent the average from two biological replicates and are shown in RPKM (Reads Per Kilobase per Million mapped reads). The dashed box highlights the 5S rDNA–SL-1 gene cluster. **(B)** Browser view zoomed in on the 5S rDNA–SL1 gene cluster. The cluster contains more than 100 tandem repeats of 5S rDNA (yellow arrowhead) and SL1 (green arrowhead) arranged in opposite orientations. **(C)** An individual example within the 5S rDNA–SL1 gene cluster. The green and yellow arrows indicate the positions and orientations of *sls-1.11* and *rrn-4.6*, respectively. **(D)** Metagene profiles of ATTF-6, Pol II (AMA-1), and Pol III (RPC-1) ChIP-seq signals around the SL1 snRNA and 5s rRNA genes. The plot is anchored at the 5’ end of the SL1 snRNA genes and spans 500 bp upstream and downstream. Signals (RPKM) were normalized to the control by subtraction and then averaged across two biological replicates. Yellow and green arrows indicate the positions and orientations of the 5S rRNA and SL1 snRNA genes, respectively.

A browser view of a single repeat with *sls-1.11* and *rrn-4.6* showed that ATTF-6 binds to the promoter region shared by the 5S rDNA and SL1 genes, with peak enrichment nearer to the SL gene (Figure 2C). Pol II was sharply enriched at *sls-1.11* (SL1 gene), while Pol III was strongly enriched at *rrn-4.6* (5S rDNA gene) (Figure 2C). Metagene analysis centered on the SL1 TSS (Transcription Start Site) further supported such organization: ATTF-6 peaked near TSS of SL1, Pol II was highest directly over SL1, and Pol III peaked over the 5S rDNA gene body (Figure 2D). Collectively, the published ChIP-seq data (Ikegami and Lieb 2013; Wang et al. 2025) and our fluorescence microscopy data suggest that Pol II and Pol III each form nuclear foci that correspond to their respective transcriptional targets at the 5S rDNA–SL1 cluster.

### Pol II and Pol III foci are dynamic and exhibit some properties of liquid-like condensates

Previous studies suggest that Pol II and several transcription factors form liquid-like condensates, potentially through liquid–liquid phase separation (Hnisz et al. 2017; Boehning et al. 2018; Cho et al. 2018; Palacio and Taatjes 2022). We therefore asked whether Pol II and Pol III foci at the 5S rDNA–SL1 cluster display properties of biomolecular condensates. These properties include dynamic molecular exchange with the surrounding environment, sensitivity to aliphatic alcohols such as 1,6-hexanediol (although this response is not universally observed), and fusion between neighboring foci. (Brangwynne et al. 2009; Patel et al. 2015; Gao et al. 2022; Alberti et al. 2025).

We first assessed molecular dynamics using Fluorescence Recovery After Photobleaching (FRAP). One of the two ATTF-6, Pol II, or Pol III foci within a nucleus was photobleached, while the other focus was left unbleached as an internal reference (Figure 3A-3C). Fluorescence recovery was monitored over 30 seconds. Recovery curves for Pol II and Pol III were well described by a single-exponential model, while ATTF-6 recovery could not be fitted by this model (Figures 3D). We found that Pol II and Pol III foci displayed much faster and more complete recovery than ATTF-6 foci (Figures 3A-3E). Specifically, the half-time of recovery (t_1/2_) was 3.41 seconds for Pol II, and 4.35 seconds for Pol III (Figure 3E). Thus, Pol II and Pol III foci exhibit rapid molecular exchange with the surrounding nucleoplasm when compared to ATTF-6.

**Figure 3.**
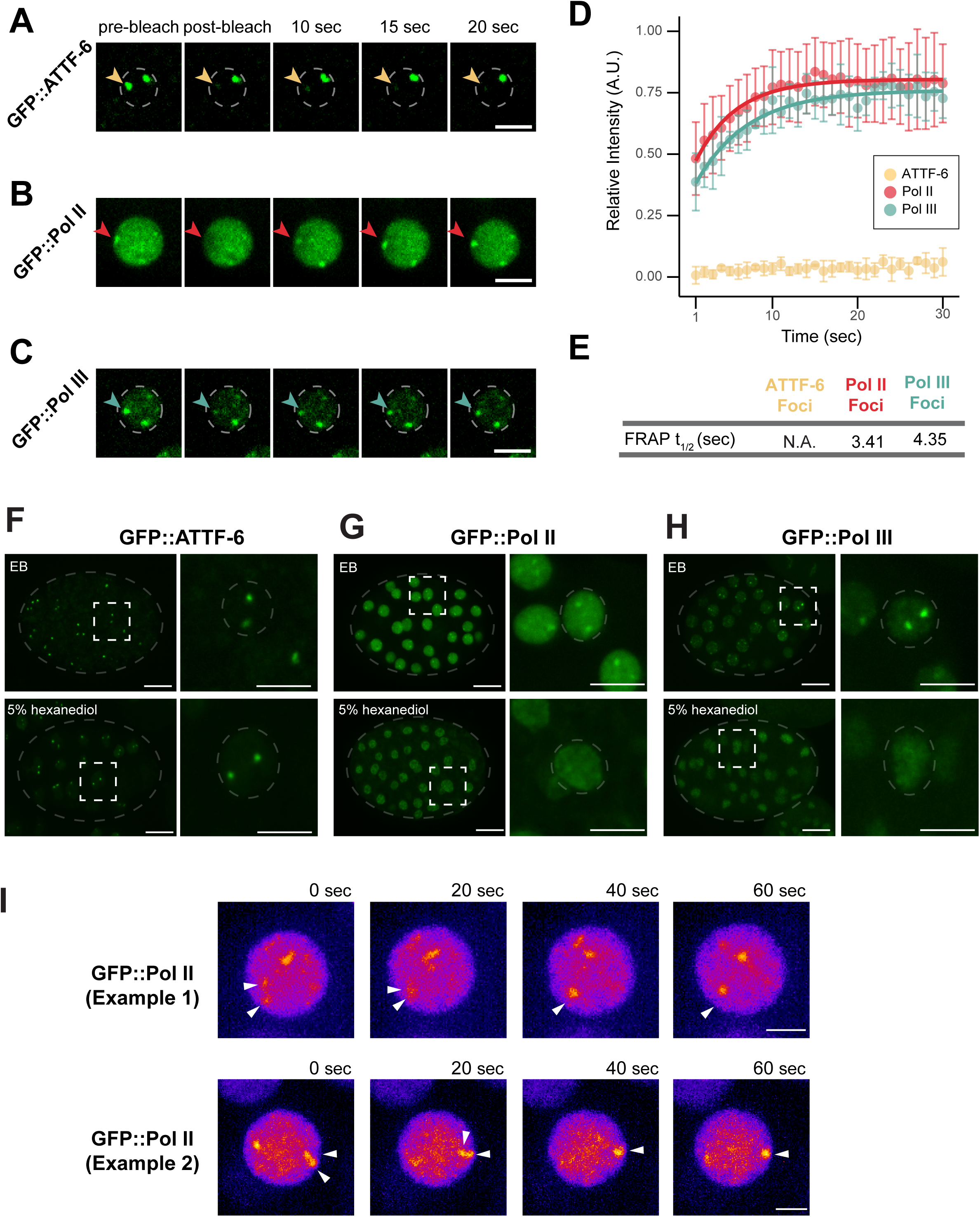
RNA Pol II and Pol III form dynamic foci *in vivo*. **(A)** Single-plane time-lapse (100x) images of GFP::ATTF-6 before (pre-bleach) and after photobleaching for the indicated timepoints (seconds). Yellow arrow indicates the photobleached focus; dashed circle indicates the nuclear periphery. Scale bar: 5 µm. **(B)** Same as in (**A**) but for GFP::Pol II. Red arrow indicates the photobleached focus. Scale bar: 5 µm. **(C)** Same as in (**A**) but for GFP::Pol III. Blue arrow indicates the photobleached focus; dashed circle indicates the nuclear periphery. Scale bar: 5 µm. **(D)** Relative intensity of GFP::ATTF-6 (n=4), GFP::Pol II (n=6) and GFP::Pol III (n=4) foci, respectively, after photobleaching in embryos. Intensity was measured from single-plane images every second (sec) before and after photobleaching and normalized to the mean intensity over five timepoints prior to bleaching (time 0). A.U. indicates arbitrary units. Dots represent mean relative intensity, and error bars denote standard deviation. Red and blue curves represent the exponential model of recovery for Pol II or Pol III, respectively. **(E)** Summary of fluorescence recovery after photobleaching (FRAP) parameters for GFP::ATTF-6, GFP::Pol II, and GFP::Pol III foci. Recovery half-times (t_1/2_) for GFP::Pol II and GFP::Pol III were calculated from the single-exponential fits shown in **(D)**. The t_1/2_ for ATTF-6 foci are labeled as N.A. (Not Applicable) because the recovery did not fit the exponential model. **(F-H)** Representative maximum intensity projections (60x) of GFP::ATTF-6 (**F**), GFP::Pol II (**G**), or GFP::Pol III (**H**) expressing embryos treated with *ptr-2* RNAi and subsequently dissected into either egg buffer (EB) or 5% 1,6-hexanediol in the EB. Images depict whole embryos, with each embryo outlined by grey dashed lines (left; scale bar: 10 µm) and the corresponding inset indicated by the white dashed square (right; inset scale bar: 5 µm). Grey dashed circles in each inset outline the nucleus. **(I)** Time-lapse fluorescence micrographs showing two collision and fusion events involving GFP::Pol II foci in early embryos. Two fusion events were observed among 135 nuclei examined. Coalescing foci are indicated by white arrowheads. Scale bar: 2 µm.

As a separate measure of focus stability, we examined ATTF-6 and polymerase foci following embryo extrusion. Embryos were extruded by gently compressing them between a microscope slide and coverslip. Following extrusion, individual nuclei were identified using fluorescently tagged histone H2B::mCherry. GFP::ATTF-6 foci remained detectable for at least 12 minutes after extrusion (Figures S3A and S3B). In contrast, GFP::Pol II foci were no longer detectable at the first imaging time point (Figures S3C and S3D), while GFP::Pol III foci were reduced at the first imaging time point and declined further over 12 minutes (Figures S3E and S3F). These data indicate that Pol II foci are less persistent than ATTF-6 and Pol III foci following mechanical extrusion from embryos.

Because 1,6-hexanediol disrupts certain types of liquid-like condensates by interfering with weak hydrophobic interactions (Alberti et al. 2025; Zheng et al. 2025), we tested the effect of 1,6-hexanediol on ATTF-6, Pol II and Pol III foci. 5% 1,6-hexanediol treatment did not dissolve ATTF-6 foci (Figure 3F). In contrast, Pol II and Pol III foci were disrupted under the same treatment conditions (Figures 3G and 3H). Thus, the polymerase foci are more sensitive to 1,6-hexanediol than ATTF-6 foci at the 5S rDNA– SL1 locus.

To determine whether these foci undergo fusion, we monitored their dynamics in early embryos using time-lapse microscopy. We examined 152 nuclei expressing GFP::ATTF-6 and 161 nuclei expressing GFP::Pol III over approximately 14 minutes but did not detect fusion events. In contrast, among 135 nuclei expressing GFP::Pol II, we observed two events in which initially separate foci merged into a single focus within 1 minute (Figure 3I). These findings suggest that Pol II foci can undergo fusion, although such events are rare under the conditions examined. This low frequency may reflect their association with chromosomal loci, which may restrict focus mobility and opportunities for contact.

In summary, these results distinguish relatively stable ATTF-6 foci from more dynamic Pol II and Pol III assemblies. Pol II and Pol III foci exhibit rapid molecular exchange, sensitivity to 1,6-hexanediol, and limited stability following embryo extrusion. Pol II foci also undergo rare fusion events. We conclude that these features are consistent with condensate-like behavior, although they do not establish that the polymerase foci form through liquid–liquid phase separation or chromatin-scaffolded assembly.

### Formation of Pol II and Pol III foci is regulated across the cell cycle

Because Pol II and Pol III foci were not detected in every nucleus within the same embryo (Figures 1 and 3), we asked whether this variability reflected differences in cell-cycle stage. Early *C. elegans* embryonic divisions consist only of S phase and M phase (Kipreos and van den Heuvel 2019). We used H2B::mCherry, which appears diffuse within the nucleus during S phase and becomes bright and highly condensed as chromosomes entering mitosis, to monitor cell-cycle stages (Figures 4A and 4B).

**Figure 4:**
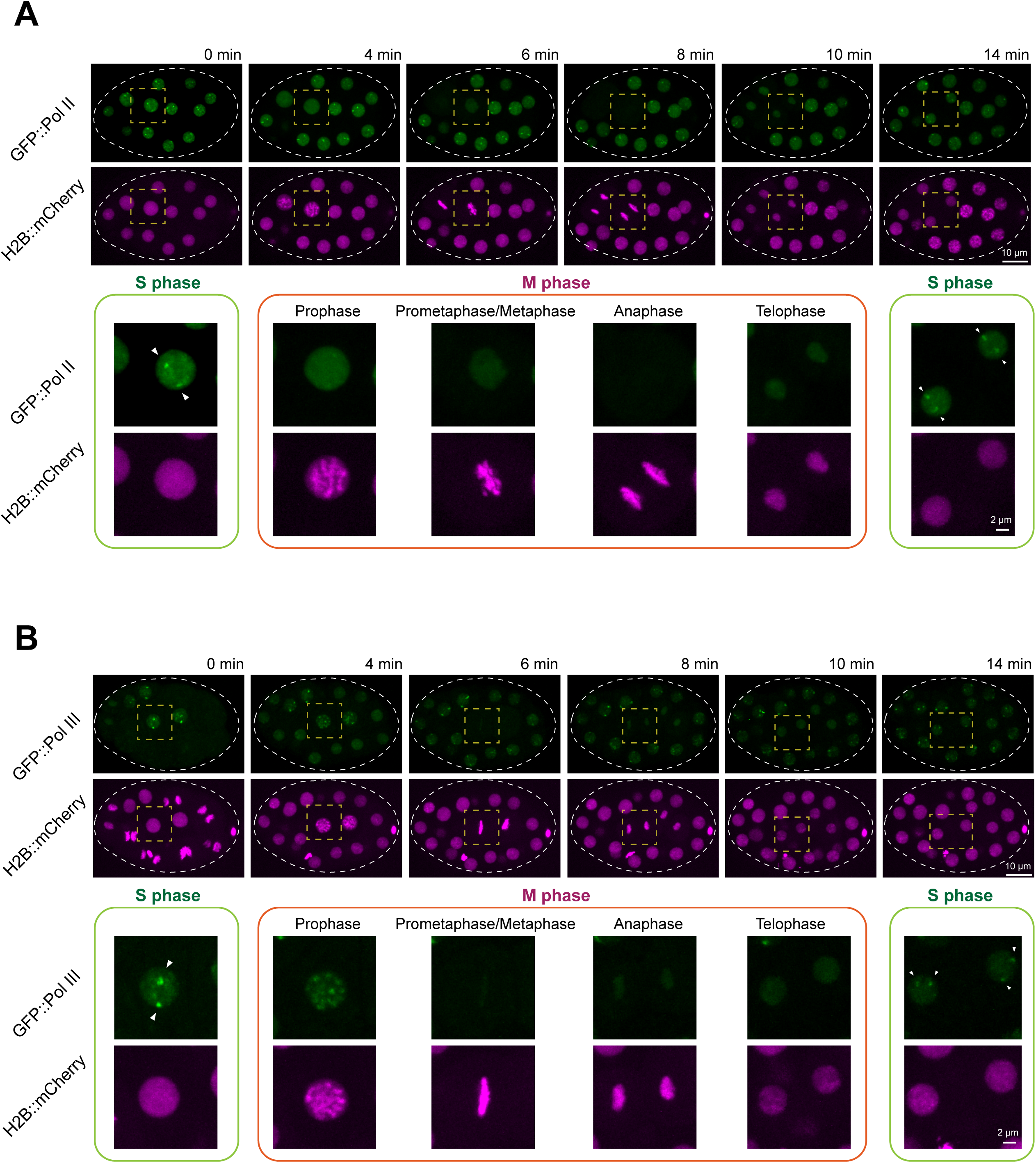
Formation of Pol II and Pol III foci is regulated across the cell cycle. **(A)** Dynamics of GFP::Pol II foci during cell division in an early embryo. A time-lapse imaging series of an embryo expressing GFP::Pol II and the chromatin marker H2B::mCherry. The circular dashed line outlines the embryo, and the dashed square highlights the dividing cell. The bottom panel shows zoomed-in views of the dividing cell, grouped into S phase and M phase. Arrowheads indicate GFP::Pol II foci in the nucleus of the dividing cell. Confocal images (60× objective) are shown as maximum-intensity projections spanning the top layer of nuclei in the embryo. Scale bars: 10 μm (top panel) and 2 μm (bottom panel). **(B)** Same analysis as in (**A**), but with GFP::Pol III.

Using this H2B::mCherry marker, we followed the dynamics of Pol II and Pol III foci within a single embryonic cell cycle. Pol II foci were consistently observed during S phase (Figure 4A). As the cell entered prophase, Pol II foci diminished in intensity and fully dispersed during mitosis. After division, Pol II foci reappeared rapidly in the S phase of each daughter cell (Figure 4A).

Pol III foci displayed a similar dependence on cell-cycle stage but with slightly distinct mitotic behavior. Pol III foci were present during S phase (Figure 4B). Upon entry into mitosis, Pol III signals redistributed onto condensed chromosomes and then became diffuse from metaphase through telophase. As with Pol II, Pol III foci re-formed in the next S phase (Figure 4B). Collectively, these observations show that Pol II and Pol III foci undergo disassembly and reassembly during the embryonic cell cycle. Both types of foci are present during S phase, disassemble as chromosomes condense during mitosis, and re-form after cell division. This reversible behavior indicates that these polymerase foci are dynamic assemblies rather than stable aggregates. Their cell-cycle-dependent organization may reflect mitotic transcriptional silencing and chromatin condensation, although the underlying mechanisms remain to be determined.

### Pol II and Pol III are dispensable for the assembly of ATTF-6 foci at the 5S rDNA– SL1 locus

Because ATTF-6 colocalizes with both Pol II and Pol III at the 5S rDNA–SL1 cluster (Figure 1), we asked whether the formation of ATTF-6 foci and polymerase foci are interdependent. To test whether polymerase activity acts upstream of ATTF-6, we examined ATTF-6 localization following depletion of Pol II or Pol III using RNAi (RNA interference) (Fire et al. 1998; Kamath et al. 2003). In embryos expressing GFP::ATTF-6 and RFP::Pol II from their endogenous loci, RNAi against Pol II efficiently reduced RFP::Pol II signals (Figures 5A and 5B). However, it had no effect on GFP::ATTF-6 foci (Figures 5A and 5B). Likewise, depletion of Pol III did not alter ATTF-6 foci formation (Figures 5C and 5D). These results indicate that neither Pol II nor Pol III is required for assembly of ATTF-6 foci.

**Figure 5.**
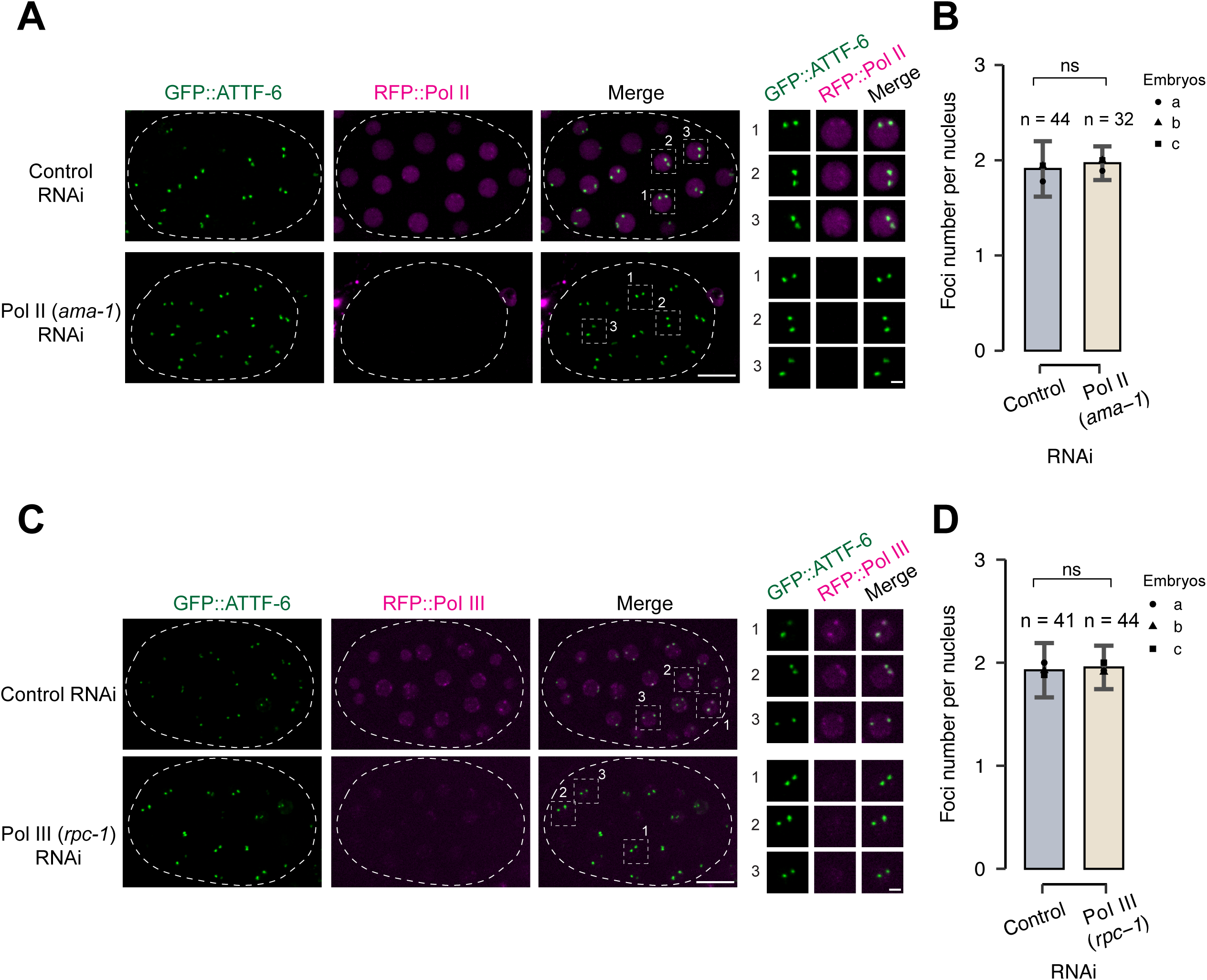
Pol II and Pol III are dispensable for assembly of ATTF-6 foci. **(A)** Fluorescence micrographs of nuclei in live embryos expressing GFP::ATTF-6 and RFP::Pol II. Embryos were dissected from worms treated with control (L4440) or Pol II (*ama-1*) RNAi. The circular dashed line outlines the embryo. Images are shown as maximum-intensity projections spanning the top nuclear layer. The right panel presents individual enlarged views of three nuclei selected from the embryo. Confocal images were acquired with a 60× objective. Scale bars: 10 μm (embryo) and 2 μm (nuclei). (B) Quantification of GFP::ATTF-6 foci numbers in embryo nuclei. Embryos were dissected from worms treated with control (L4440) or Pol II (*ama-1*) RNAi. Nuclei from three independent embryos were counted. Error bars represent the standard deviation (SD). Statistical significance was determined using a two-tailed Student’s t-test (ns: p > 0.05). **(C-D)** Analyses as in (**A-B**), but using embryos dissected from worms treated with Pol III (*rpc-1*) RNAi.

### ATTF-6 is required for Pol II foci formation but is dispensable for Pol III foci

We next performed the reciprocal test to determine whether ATTF-6 is required for polymerase foci formation. Since *attf-6* is an essential gene (Wang et al. 2025), we depleted ATTF-6 protein using the auxin-inducible degron (AID) system (Zhang et al. 2015). Using CRISPR/Cas9 gene editing, we tagged ATTF-6 at its endogenous locus with RFP and AID, enabling both visualization and auxin-dependent degradation. Germline- and embryo-specific degradation of ATTF-6 was achieved using a *sun-1p::TIR1* transgene, in which the *sun-1* promoter drives expression of TIR1 (an auxin-dependent E3 ubiquitin ligase receptor) in the germline and embryos (Ashley et al. 2021). We performed live imaging of embryos expressing ATTF-6::AID::RFP together with GFP::Pol II or GFP::Pol III (Figure 6A), and quantified both the number of foci per nucleus (Figure S4) and the fraction of nuclear fluorescence localized within foci (Figures S5).

**Figure 6.**
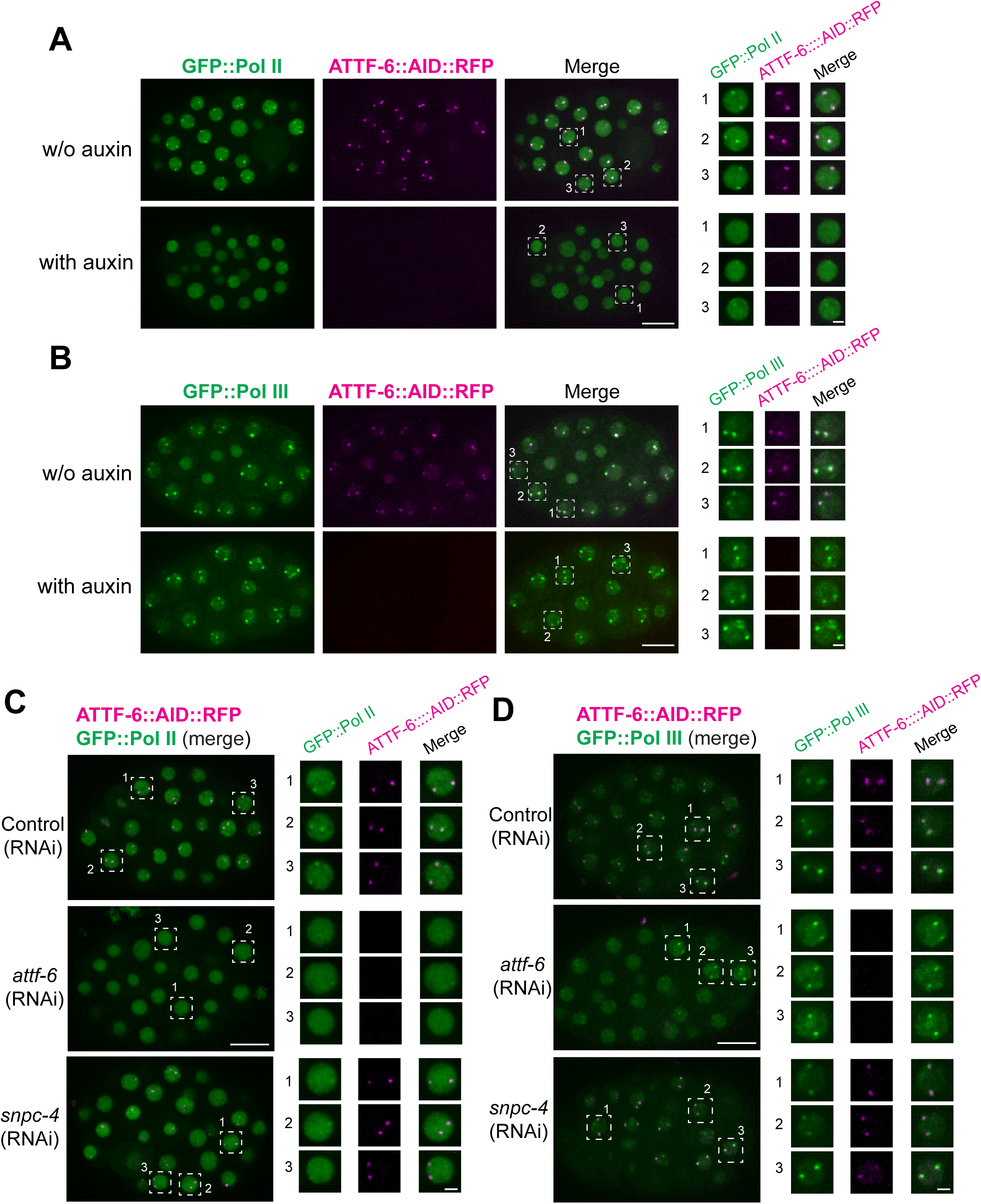
SNPC-4 and ATTF-6 are required for Pol II foci, but not Pol III foci formation. **(A)** Live embryos expressing GFP::Pol II and ATTF-6::AID::RFP, dissected from worms treated with or without 1 mM auxin. The right panel presents individual enlarged views of three nuclei selected from each embryo. Confocal images (60× objective) are shown as maximum-intensity projections spanning the top nuclear layer. Scale bars: 10 μm (embryo) and 2 μm (nuclei). **(B)** Same auxin treatment conditions as in **(A),** but using embryos expressing GFP::Pol III and ATTF-6::AID::RFP. **(C)** Live embryos expressing GFP::Pol II and ATTF-6::AID::RFP, dissected from worms treated with control (L4440), *snpc-4*, or *attf-6* RNAi. Embryos show the merged GFP and RFP signals. The right panels present individual enlarged views of three nuclei selected from each embryo, shown as separate GFP, RFP, and merged channels. Confocal images (60× objective) are shown as maximum-intensity projections spanning the top nuclear layer. Scale bars: 10 μm (embryo) and 2 μm (nuclei). **(D)** Same RNAi conditions as in (**C**), but showing embryos expressing GFP::Pol III together with ATTF-6::AID::RFP.

In the absence of auxin, GFP::Pol II formed two prominent nuclear foci that colocalized with ATTF-6::AID::RFP (Figures 6A). Upon auxin treatment, ATTF-6::AID::RFP foci became undetectable (Figures 6A, S4A, S4B, S5A and S5B). Depletion of ATTF-6 resulted in the loss of large Pol II foci (Figures 6A, S4A and S4B). The fraction of nuclear GFP::Pol II fluorescence localized within foci was reduced from 11.9% in the untreated control to 1.3% in the auxin-treated sample (Figure S5A). In contrast, depletion of ATTF-6 only modestly altered the number of GFP::Pol III foci (Figures 6B, S4A, and S4C), and the fraction of nuclear GFP::Pol III fluorescence localized within foci was modestly reduced (Figure S5B). These findings demonstrate that ATTF-6 is required for Pol II focus formation but is largely dispensable for Pol III focus formation.

To independently confirm this requirement and to identify additional factors involved in polymerase foci assembly, we used RNAi to deplete *attf-6* as well as *snpc-4*, a subunit of Small Nuclear RNA-activating Protein Complex that drives the transcription of snRNA genes (Kasper et al. 2014; Hou et al. 2022). Consistent with the phenotype observed following AID-mediated depletion of ATTF-6 protein, RNAi-mediated knockdown of *attf-6* caused dispersal of Pol II foci (Figures 6C, S4D and S5C), without substantially affecting Pol III focus number or the fraction of nuclear Pol III fluorescence localized within foci (Figures 6D, S4E, and S5D). Knockdown of *snpc-4* did not disrupt ATTF-6 localization. However, *snpc-4* knockdown impaired GFP::Pol II focus formation, while GFP::Pol III foci remained detectable (Figures 6D, S4E, and S5D). Taken together, these results show that ATTF-6 and SNPC-4 are specifically required for the formation of Pol II foci at the 5S rDNA–SL1 cluster, while Pol III foci assemble independently of these factors.

### Pol II foci are temperature-sensitive and regulate SL1 expression

To investigate the functional significance of polymerase foci in regulating SL1 and 5S rRNA expression, we examined how their assembly responds to environmental cues. *C. elegans* is a poikilothermic organism whose cellular processes adapt to environmental temperature. And biomolecular condensates are often sensitive to thermal perturbation (Fritsch et al. 2021; Alberti et al. 2025). We therefore tested whether elevated temperature affects the formation of ATTF-6, Pol II, or Pol III foci.

Embryos were shifted from 20 °C to 32 °C, and imaged after 2 and 3 hours. GFP::ATTF-6 foci remained intact under these heat stress conditions (Figures 7A, S6A and S6B). In contrast, GFP::Pol II foci progressively disassembled at 32°C (Figure 7B). After 3 hours, most embryonic nuclei lacked detectable Pol II foci, and the fraction of nuclear GFP::Pol II fluorescence localized within foci was reduced by 55.5-fold relative to the 20°C control (Figures S6C and S6D). By comparison, both the number of GFP::Pol III foci and the fraction of nuclear GFP::Pol III fluorescence localized within foci remained largely unchanged at 32°C (Figures 7C, S6E, and S6F). These observations indicate that Pol II foci are more sensitive to elevated temperature than ATTF-6 and Pol III foci under the conditions examined.

**Figure 7.**
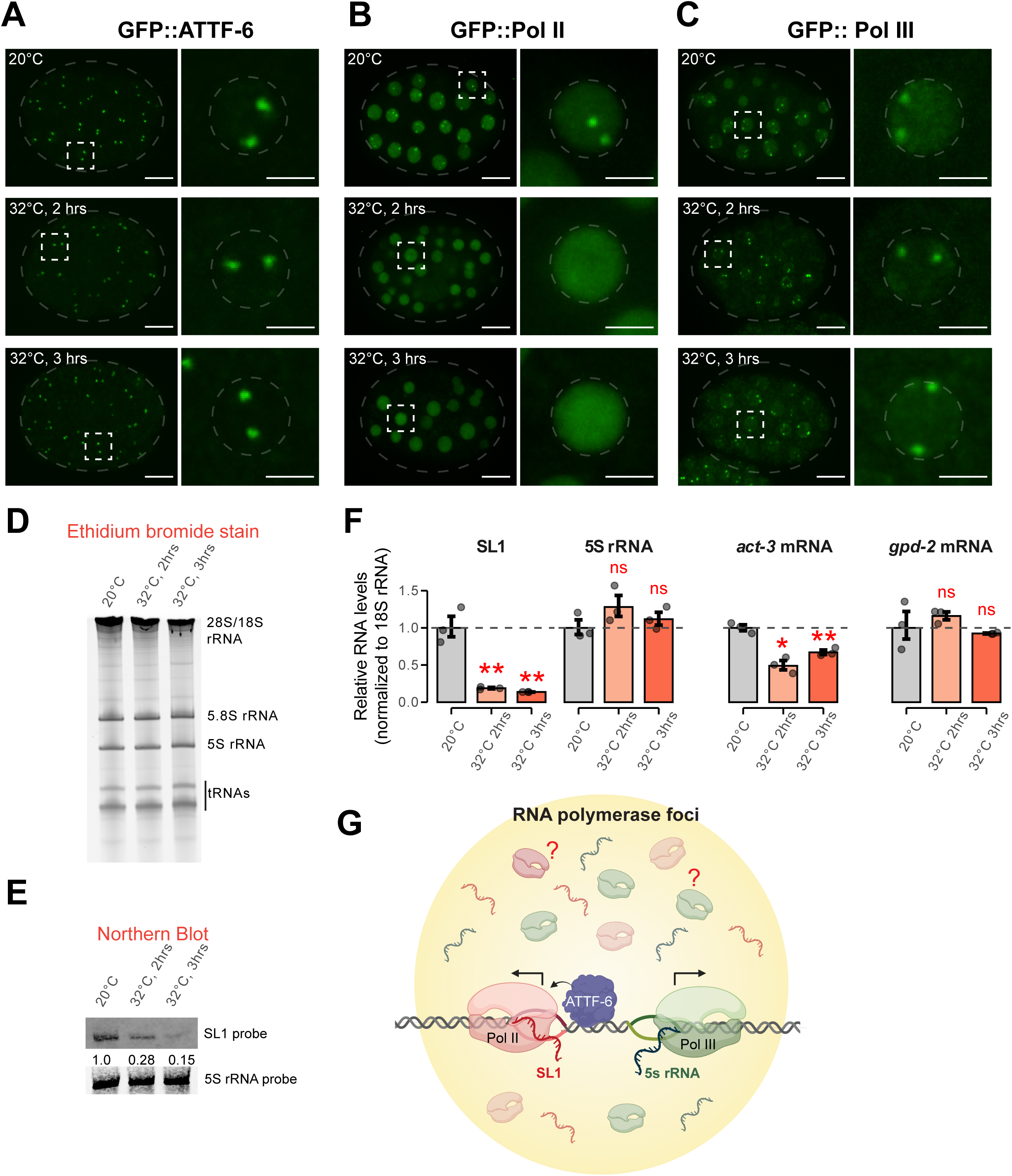
Pol II focus disruption at elevated temperature correlates with reduced SL1 expression. **(A)** Representative maximum intensity projections (60x) of GFP::ATTF-6 expressing embryos cultured at 20°C (top) or shifted to 32°C for either 2 hours (middle) or 3 hours (bottom). Whole embryos are shown on the left (scale bar: 10 µm), outlined by grey dashed ovals. The corresponding insets to the right are indicated by the white dashed squares. Insets to the right show single nuclei outlined by grey dashed circles (Inset scale bar: 3 µm). Heating conditions are further described in the Materials and Methods. **(B)** Same conditions as in (**A**), but for GFP::Pol II expressing embryos. **(C)** Same conditions as in (**A**), but for GFP::Pol III expressing embryos **(D)** Ethidium bromide staining of total RNAs from wild-type embryos cultured at 20°C, or shifted to 32°C for either 2 hours or 3 hours in M9 buffer, respectively. rRNAs and tRNAs are indicated. **(E)** Northern blotting of SL1 RNA (top) or 5S rRNA (bottom) from wild-type embryos cultured at 20°C, or shifted to 32°C for either 2 hours or 3 hours, respectively. Relative SL1 abundance upon heating was normalized to 5S rRNA abundance. **(F)** RT-qPCR analysis of Pol II transcripts (SL1, *act-3,* and *gpd-2)* and the Pol III transcript 5S rRNA in embryos. Transcript fold change upon heat stress was calculated relative to 20°C and normalized to 18S rRNA as the internal control. Bar plots show mean +/− SD; dots represent technical replicates. Statistical significance was assessed using Student’s two-sample t-test. * indicates p ≤0.05, ** p ≤0.01, and ns indicates not significant. **(G)** Working model showing Pol II and Pol III occupy the 5S rDNA–SL1 locus and assemble into nuclear compartments. ATTF-6 promotes Pol II foci formation to drive SL1 transcription, while Pol III independently associates with 5S rRNA genes.

We next investigated whether temperature-dependent changes in polymerase foci correlate with altered SL1 or 5S rRNA expression. Total RNA extracted from embryos incubated at 20 °C or 32 °C was analyzed by denaturing gel electrophoresis. Ethidium bromide staining revealed no detectable change in the abundance of major RNA species including tRNAs, 5S, 5.8S, 18S, and 28S rRNAs (Figure 7D). Northern blotting of the same samples demonstrated that SL1 RNA levels decreased ∼4-fold after 2 h at 32 °C and ∼5-fold after 3 h, when normalized to 5S rRNA signals (Figure 7E). RT–qPCR analysis corroborated these results: SL1 transcripts were significantly reduced under the heat stress, while 5S rRNA levels were unaffected when normalized to 18S rRNA (Figure 7F). Among housekeeping controls, *act-3* mRNA showed a modest decrease, while *gpd-2* levels were unchanged (Figure 7F). Together, these findings indicate that elevated temperature selectively disrupts Pol II foci and that this disassembly correlates with a marked reduction in SL1 transcription.

## Discussion

Although the spatial organization of transcription machinery is increasingly recognized as a key regulatory layer, a major knowledge gap remains: it is unclear whether different RNA polymerases form compartments when operating within the same gene cluster, and how such organization influences transcriptional output. In this study, we uncover a specialized nuclear microenvironment at the *C. elegans* 5S rDNA–SL1 cluster in which RNA polymerases II and III form dynamic compartment. Our findings support a model in which transcription factors including ATTF-6 and SNPC-4 recruit or stabilize Pol II at the SL1 gene cluster to promote efficient SL1 transcription, while Pol III independently assembles on adjacent 5S rRNA genes (Figure 7G). These two polymerases therefore occupy the same genomic locus but form functionally distinct nuclear foci.

The precise mechanism underlying the formation of polymerase foci remains unresolved. Tandem repeats at the 5S rDNA–SL1 locus provide spatially clustered binding sites for transcription factors and polymerases. This allows transcriptional machinery to become locally enriched through chromatin-scaffolded recruitment without requiring the formation of a separate liquid phase (Erdel and Rippe 2018; Muzzopappa et al. 2022). In this model, ATTF-6 functions as a stable chromatin-associated component that promotes Pol II recruitment or retention at the locus (Figure 7G). It is worth noting chromatin-scaffolded recruitment and phase separation are not mutually exclusive. Multivalent interactions among polymerases, transcription factors, chromatin, and nascent RNA further concentrate and stabilize these assemblies and potentially contribute to their condensate-like properties.

Another unresolved question is the molecular composition of these foci. Although ChIP-seq data demonstrate the occupancy of polymerases at the 5S rDNA–SL1 cluster (Figure 2), our imaging experiments do not distinguish transcriptionally engaged polymerases from unbound or transiently associated polymerases (Figure 7G). Interestingly, the C-terminal domain of Pol II can undergo phase separation in vitro and promote Pol II clustering in cells (Boehning et al. 2018), raising the possibility that a pool of unbound Pol II may contribute to focus formation. Future studies defining the molecular composition of Pol II and Pol III foci will provide further insight into how transcription is organized within repetitive gene clusters.

An emerging theme in gene regulation is that transcription can be organized into spatially restricted structures, ranging from transient hubs of polymerase-transcription factor interactions to fully formed biomolecular condensates generated by phase separation (Banani et al. 2017; Hnisz et al. 2017; Rippe 2022). Our data place Pol II and Pol III foci at the 5S rDNA–SL1 cluster closer to the latter category. Both foci display rapid fluorescence recovery after photobleaching. Furthermore, Pol II foci are sensitive to 1,6-hexanediol and thermal perturbation. These features match the operational hallmarks of liquid-like condensates. However, these polymerase foci differ from previously reported Pol II condensates (Cho et al. 2018). They are unusually large (> 600 nm), highly localized to a specific gene cluster, and assemble in a cell-cycle– dependent manner strictly during S phase. This suggests that the polymerase bodies at the 5S rDNA–SL1 locus represent a specialized class of “gene-specific compartment” tuned to the unique regulatory needs of these small non-coding RNA genes.

Although Pol II and Pol III foci occupy the same genomic region, their formation is mechanistically separable. Pol III foci assemble independently of ATTF-6 and SNPC-4 and are more stable under heat stress. Pol II foci, in contrast, require ATTF-6 and SNPC-4 and are sensitive to elevated temperature. These differences reveal distinct regulatory architectures for the two polymerases. One possibility is that Pol III foci depend more strongly on chromatin-scaffolded recruitment at the repetitive 5S rDNA locus, while Pol II foci may involve a greater contribution from multivalent interactions. However, the current genetic and imaging data do not distinguish these mechanisms. It is also important to note that elevated temperature triggers heat-shock response in *C. elegans*, including HSF-1 activation and widespread transcriptional reprogramming (Morton and Lamitina 2013; Brunquell et al. 2016; Garrigues et al. 2019). We therefore cannot exclude the possibility that the disruption of Pol II foci and the accompanying reduction in SL1 transcripts arise from broader stress-induced changes in transcriptional regulation rather than a direct effect on the stability of Pol II foci. Nevertheless, the temperature sensitivity of Pol II foci is particularly interesting, because this may position the Pol II foci as a temperature-responsive regulatory element, potentially enabling rapid transcriptional adaptation without global repression of rRNA production.

Some other clustered loci may also serve as sites of polymerase accumulation. For example, in *Drosophila*, replication-dependent histone genes are associated with the histone locus bodies, specialized nuclear compartments enriched for Pol II and factors required for histone gene processing (Huang et al. 2021; Kemp et al. 2021). Unlike *Drosophila*, *C. elegans* histone genes are organized in multiple small clusters dispersed throughout the genome and do not appear to form canonical the histone locus bodies (Duronio and Marzluff 2017). Nevertheless, our ChIP-seq analysis revealed strong Pol II enrichment at histone gene clusters in addition to the 5S rDNA–SL1 cluster (Figures 2 and S2). It is therefore possible that some of the smaller Pol II foci observed in nuclei correspond to histone gene clusters (Figure 1), although this remains to be tested experimentally.

Our results extend our previous work showing that ATTF-6 is required for SL1 accumulation but is dispensable for 5S rRNA biogenesis (Wang et al. 2025). Here we demonstrate that ATTF-6 is required for the assembly of Pol II foci at the 5S rDNA–SL1 locus, providing a plausible explanation for its selective effect on SL1 transcription. By contrast, Pol III foci remain largely intact without ATTF-6 (Figures 6 and 7), consistent with the observation that 5S rRNA levels are largely unchanged following ATTF-6 depletion (Wang et al. 2025). Although our analyses show a clear correlation between Pol II foci disassembly and reduced SL1 transcript levels, caveats remain. For example, SL1 and 5S RNA populations may differ in stability. Hence, unchanged 5S abundance under heat stress does not necessarily reflect maintained transcription. Future approaches that directly measure nascent RNA synthesis, such as metabolic labeling, will be important for resolving the transcriptional responses of both loci.

In summary, our findings support a model in which ATTF-6–dependent Pol II foci drive efficient SL1 transcription, while Pol III independently forms its own foci to produce 5S rRNA (Figure 7G). The two polymerases coexist within a shared genomic compartment yet operate through different assembly principles and heat sensitivities. This organizational strategy may allow animals to tightly couple Pol II–driven SL1 production to physiological conditions while preserving robust Pol III–mediated 5S rRNA synthesis.

## Acknowledgement

We thank members in the Tang lab for discussion and critical comments, OSU Neuroscience Imaging Core for instruments (S10OD026842), the Caenorhabditis Genetics Center for providing the *C. elegans* strains (P40OD010440). Y.H. Wang was supported by Center for RNA Biology graduate fellowship and presidential fellowship at The Ohio State University. This work was supported by the Arthur Burghes professorship, NIH Maximizing Investigators’ Research Award (R35 GM142580), and NSF (MCB 2420329) to W. Tang.

## Author contributions

W.T., YH.W. and H.L.H. conceptualized the study. YH.W., H.L.H. and W.T. performed experiments and analyzed the data. W.T. supervised the study and acquired funding. YH.W., H.L.H. and W.T. wrote the manuscript.

## Data availability

ATTF-6 ChIP-seq data were previously published and are available at NCBI under the accession number GSE277641 (Wang et al. 2025). Pol II and Pol III ChIP-seq data were previously published and are available at NCBI under the accession number GSE42741 (Ikegami and Lieb 2013).

## Materials and Methods

### Maintenance of *C. elegans* strains

All strains were grown at 20°C unless otherwise stated. Wild-type refers to the N2 Bristol strain (Brenner 1974). GFP::AMA-1 and GFP::RPC-1 strains were obtained from the CGC and are referred to as GFP::POL II and GFP::POL III, respectively, for clarity throughout this manuscript. A complete strain list may be found in Supplementary Table S1.

### CRISPR/Cas9 genome editing

CRISPR/Cas9 genome editing in *C. elegans* was carried out as previously described (Kim et al. 2014). The ATTF-6::AID::RFP strain was generated using AID::RFP double-stranded DNA donors in a *sun-1p*::TIR1; *gfp::ama-1* background. The RFP::Pol II and RFP::Pol III strains were produced using RFP double-stranded DNA donors in the GFP::ATTF-6 background. Donor DNA was mixed with a pre-assembled Cas9 ribonucleoprotein complex consisting of Cas9 protein, gRNA, and tracrRNA (IDT). The pRF4 plasmid carrying the dominant rol-6 allele was included as a co-injection marker (Kim et al. 2014). F1 roller progeny were selected and screened by PCR. Homozygous insertion strains were confirmed by Sanger sequencing. DNA donor templates, gRNA sequences, and genotyping primers are listed in Supplementary Table S2.

### Embryo extrusion

For ex vivo extrusion experiments, GFP::ATTF-6, GFP::Pol II, or GFP::Pol III adult gravid hermaphrodites were treated with bleaching solution containing 1.25% sodium hypochlorite and 0.25 M NaOH for 10 minutes to isolate embryos and weaken the eggshell. Bleached embryos were washed three times in Egg buffer (or EB: 25 mM HEPES-NaOH, pH 7.4, 118 mM NaCl, 48 mM KCl, 2 mM EDTA, 0.5 mM EGTA) and then transferred to 4% agar slides. To mechanically extrude embryonic nuclei, two diagonal corners of the cover slip were firmly compressed, and immediately imaged (<5 minutes after extrusion). Time zero was defined as the first image acquired following extrusion. Images were acquired every 3 mins (GFP::ATTF-6, GFP::Pol III) or 4 mins (GFP::Pol II), respectively for a 12 minute imaging period.

### DNA-FISH and immunofluorescence

The approach was adapted from an established protocol with modifications (Phillips et al. 2009; Adilardi and Dernburg 2022). Briefly, ∼5000 gravid adult worms (ATTF-6::3×FLAG) were bleached and washed with Egg Buffer. Approximately 50 eggs in 20 μL EB were seeded onto a coverslip and fixed by adding 20 μL of 4% formaldehyde in EB to reach a final concentration of 2%, followed by incubation for 2–3 min. After fixation, the eggs were covered with a poly-L-lysine–coated slide. The slide was placed on a metal block pre-cooled with dry ice for 30 min. The coverslip was then carefully flicked off, and the slide was immediately immersed in −20°C methanol for 10 min. Residual liquid was removed from around the eggs, and the samples were washed once with 2× SSCT (0.3 M NaCl, 0.03 M sodium citrate, pH 7.0, 0.1% Tween-20). The eggs were then incubated in 2× SSCT containing 50% formamide at 37°C overnight. For DNA probe hybridization, synthesized Cy5-labeled DNA probes (10 ng/μL; IDT) were diluted in hybridization buffer (HB: 3× SSC, 48% formamide, 10.6% dextran sulfate). The probe mixture was added to the samples, covered with a coverslip, and heated at 80°C for 10 min. Slides were then transferred to a light-protected humid chamber and incubated at 37°C for 6 h. After hybridization, egg samples were washed three times with PBST (PBS + 0.1% Tween-20) and incubated in blocking buffer (PBST containing 0.5% BSA) at room temperature for 1 h. Mouse anti-FLAG primary antibodies (Sigma-Aldrich) were diluted 1:500 in blocking buffer and applied to the samples overnight at 4°C. After three PBST washes, samples were incubated with goat anti-mouse Alexa Fluor 488 secondary antibody (1:500 in blocking buffer) for 2 h at room temperature in the dark. Following incubation, samples were washed three times with PBST. A drop (10 μL) of DAPI-containing mounting medium (Vector Laboratories) was applied to the eggs and incubated for 10 min at room temperature. Samples were then covered with a glass coverslip and sealed with nail polish.

### RNA interference by feeding of double stranded RNA

HT115 RNAi feeding bacteria were streaked from the Ahringer RNAi library as described previously (Kamath et al. 2003; Price et al. 2021). In brief, Nematode Growth Medium (NGM) plates containing 50 µg/mL ampicillin and 5 mM IPTG were seeded with HT115 bacteria expressing double-stranded RNA (dsRNA) targeting the gene of interest. L1 larvae were transferred onto RNAi plates and grown at 25°C for two days. Gravid worms were then dissected, and their embryos were imaged. L4440 RNAi was used as the control for all RNAi experiments.

### Auxin treatment

The auxin treatment was performed as previously described (Zhang et al. 2015; Wang et al. 2025). Briefly, *attf-6::aid::3xflag::rfp*; *sun-1p::TIR1* reporter strains (Table S1) were transferred to NGM plates containing 1 mM auxin indole-3-acetic acid (IAA) at the L3– L4 stage and incubated at 25°C overnight. Adult worms were imaged to confirm complete depletion of ATTF-6::AID::3xFlag::RFP in the germline and embryo. Subsequently, adult worms were dissected, and the resulting embryos were imaged for further analysis.

### Granule disruption with hexanediol or heat stress

For 1,6-hexanediol (Sigma-Aldrich) treatment, L4 worms expressing GFP::ATTF-6, GFP::Pol II, or GFP::Pol III, respectively, were plated on *ptr-2* RNAi plates to permeabilize the eggshell of embryos for 1,6-hexanediol treatment and fed for approximately 24 hours (Thomas et al. 2023). At adulthood, *ptr-2* RNAi treated adults were dissected into either egg buffer (EB) or 5% 1,6-hexanediol in EB on ring slides and imaged.

To disrupt granules via heating, GFP::ATTF-6, GFP::Pol II, or GFP::Pol III expressing gravid adults grown at 20°C were bleached to collect embryos. For the control condition (20°C), resuspended embryos were mounted on glass slides with 4% agarose pads and imaged immediately. For heating, embryos were resuspended in ∼ 4 mL M9 in a 15 mL conical tube and transferred to a 32°C incubator (VWR Scientific Model 1545) shaking on a nutating mixer (Fisher Scientific). Embryos were imaged at ambient temperature either 2 hours or 3 hours post-transfer to 32 °C.

### Heat-stress conditions and RNA extraction

Wild-type gravid adults grown at 20°C were bleached to obtain embryos. For each condition, 600,000 embryos were resuspended with 9 mL M9 in 15 mL conical tubes. For the control condition (20°C), RNA was extracted from resuspended embryos following bleaching. For the heat-stress conditions, RNA was extracted from resuspended embryos transferred to 32°C (VWR Scientific Model 1545) and incubated on a nutating mixer (Fisher Scientific) for either 2 hours (32°C, 2 hours) or 3 hours (32°C, 3 hours). For all three conditions, respectively, samples were pelleted and resuspended in 1ml of TRI Reagent (Sigma-Aldrich).

Total RNA was extracted as follows. Samples were lysed using the Bead Mill 24 homogenizer (Thermo Fisher Scientific) and the lysate was transferred to 1.5 ml centrifuge tube. 0.1 volumes of bromochloropropane (Sigma-Aldrich) were added to the mix and shaken for 30 seconds, then centrifuged at 15,000 × g. The aqueous phase was transferred to a fresh 1.5 mL tube, and 1 volume of isopropanol was added. Samples were incubated at −20°C for 1 hour to precipitate RNA, followed by centrifugation at 15,000 × g for 15 minutes at 4°C. The supernatant was removed, and the RNA pellet was washed twice with 1 mL 75% ethanol. Pellets were air dried for 3 minutes, dissolved in ddH₂O, and stored at −80°C.

### Ethidium bromide staining

To assess overall RNA abundance, 7.5 µg of total RNA was separated on an 8% polyacrylamide gel containing 7 M urea. The gel was stained with 5 µg/mL ethidium bromide (Sigma-Aldrich) for 15 min, washed twice with 1× TBE buffer, and imaged using a GelDoc Gel imaging system (Bio-Rad).

### Northern Blotting

To quantify SL1 or 5s rRNA abundance, 15 µg total RNAs were separated on an 8% polyacrylamide/7M urea gel and transferred to Hybond N+ membrane (GE healthcare) at 400 mA for 1 hour with 1x TBE buffer. RNAs were crosslinked to the membrane using ultraviolet light (254 nm; 120 mJ) with a Stratalinker (Stratagene). Membranes were prehybridized with Ultrahyb Ultrasensitive Hybridization Buffer (Invitrogen). Subsequently, 10 µmol of either fluorescein amidite (FAM)-labeled anti-SL1 or anti-5s rRNA probes were hybridized to their respective membranes at 37 °C, rotating overnight. The following day, the membranes were washed three times with wash buffer (0.1% sodium dodecyl sulfate, 0.1% saline-sodium citrate) at 37 °C and visualized using a Sapphire Biomolecular Imager (Azure Biosystems). SL1 and 5S rRNA abundance was quantified using FIJI.

### RT-qPCR (Real-time quantitative PCR)

100 ng of total RNAs from embryos were reverse transcribed via Multiscribe Reverse Transcriptase (Thermo Fisher Scientific) using gene-specific antisense RT primers containing a universal stem loop. Real-time quantitative PCR was conducted with diluted RT reactions (1/4 for SL1, *act-3*, and *gpd-2*; 1/100 for 5S rRNA and 18S rRNA) using the respective gene-specific forward primer and an antisense primer matching the universal stem loop (Pastore et al. 2021). Reactions were conducted in triplicate using PowerUP SYBR Green Master Mix (Thermo Fisher Scientific) and run with the CFX Connect Real-Time PCR System (Bio-Rad). Transcript levels were normalized to the reference gene 18S rRNA, and relative levels upon heating was calculated via the ΔΔCt method. Primer sequences listed in Supplementary Table S2.

### Analysis of ChIP-seq data

TrimGalore was used to trim adapter sequences and filter out low-quality reads (Kechin et al. 2017). The resulting high-quality reads were aligned to the genome with the BWA-MEM algorithm under default settings (Vasimuddin et al. 2019). Following alignment, BAM files were transformed into BigWig files using DeepTools bamCoverage function (Ramírez et al. 2016). Enrichment peaks from ChIP-seq data were detected with MACS2 employing the narrow-peak option (Zhang et al. 2008; Feng et al. 2012). BigWig signals from two biological replicates were averaged using the DeepTools bigwigMerge function. To compare IP and control signals, experimental BigWig files were subtracted from control BigWig files using bigwigCompare with the parameters (-p max -operation subtract -pseudocount 1 -binSize 5). Further processing and statistical analyses were carried out with custom R and Python pipelines. Visualization of BigWig files was performed using IGV (Robinson et al. 2011). ChIP-seq metagene profiles were generated using the SeqPlots software (Stempor and Ahringer 2016). The subtracted BigWig files from the ChIP-seq pipeline served as input, and BED files containing SL1 snRNA genes were used as references for the plots.

### Microscopy

Unless otherwise indicated in the figure caption, all images depict live embryos dissected from gravid adults into EB and mounted on glass slides with 4% agarose pads.

Most imaging was conducted using the Nikon Ti2 inverted microscope equipped with an X-Light V3 spinning disk confocal unit (CrestOptics) in NIS-Elements AR 5.41.02, using the Plan Apo 60x water objective (Price et al. 2023) with one exception: to determine fluorescence recovery after photobleaching (FRAP) of ATTF-6, Pol II, and Pol III foci in embryos, spinning disk confocal images were taken using a Nikon TiE inverted microscope equipped with an Andor Revolution WD spinning disk system. Images were captured with a CFI Plan Apo VC 100x/1.4 NA oil immersion objective and Andor iXon Ultra897 camera. All further image processing was conducted with FIJI.

### Image Quantification

Photobleaching of GFP::ATTF-6, GFP::Pol II, or GFP::Pol III foci was conducted using live embryos dissected into EB. Foci were photobleached using the FRAPPA module in MetaMorph with a 70% 488 nm laser, and a dwell time <1 ms per pixel, and either a single pulse (ATTF-6) or two pulses (Pol II and Pol III) using a rectangular Region of Interest (ROI) encompassing the foci. Images were acquired every second for 5 seconds prior to and 30 seconds post photobleaching. Images were further analyzed in FIJI.

For each focus, the fluorescence intensity at each timepoint (I_t_) was background-corrected with intensity outside the embryo (I_b_). The mean pre-bleach intensity (I_0_) was calculated from 5 frames prior to bleaching, and corrected to I_b_. The normalized fluorescent intensity was then computed as: 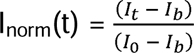.

To determine whether ATTF-6, Pol II, or Pol III exhibited measurable recovery, the maximum post-bleach intensity was compared to the baseline intensity at time zero. Recovery was considered significant if the increase in intensity exceeded twice the standard deviation of the baseline, to avoid fluctuations due to noise. For proteins that met the above criteria, the mean I_norm_ was calculated at each timepoint, and the mean value at the first post-bleach frame was defined as y_0_. FRAP recovery was modeled in R with a first-order exponential function: F(t) = y_0_ + A_rec_ * (1-e^−***k***Time^), where A_rec_ is the fluorescence recovery amplitude and ***k*** is the recovery rate constant. Curve fitting was performed using the Levenberg–Marquardt algorithm implemented in the nlsLM function from the minpack.lm R package. t_1/2_ was calculated from the first order exponential model.

To quantify the number of foci per nucleus for GFP::ATTF-6, GFP::Pol II, or GFP::Pol III embryos, maximum intensity projections spanning the top single layer of nuclei (6.6 µm, step=0.3 µm) were generated. Circular ROIs were drawn surrounding each nucleus and foci were manually counted using the Multipoint tool in FIJI. The number of points bound by each ROI was calculated in FIJI to calculate foci numbers. Three embryos were quantified per condition.

To determine the fraction of nuclear fluorescence contained within the GFP::ATTF-6, GFP::Pol II, or GFP::Pol III foci, at least three embryos were selected per condition. Maximum intensity projections were generated spanning the top 6.6 µm of each embryo (step = 0.3 µm). ROIs encompassing individual nuclei were drawn for analysis. Background fluorescence was removed using a rolling-ball subtraction radius of 140 µm. Foci were identified using the following thresholds: 20-100% of maximum pixel intensity for ATTF-6; 40-100% for Pol III; and 65-100% for Pol II. To exclude noise, only particles with an area of 0.05–1.2 µm^2^ were analyzed.

For each nucleus, the fraction of nuclear fluorescence localized within foci was calculated by dividing the summed integrated density of all detected foci by the total integrated density of the nucleus and multiplying by 100:

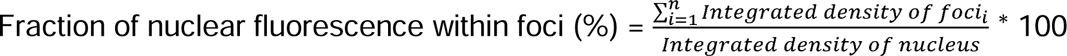

where n denotes the number of detected foci within each nucleus. When no foci were detected in the nucleus, the numerator was set to zero.

---

**Figure S1. RNA Pol II and Pol III foci are formed at the 5S rDNA–SL1 locus in the germline.**

**(A)** Schematic of the *C. elegans* germline and embryos in utero. Colored lines highlight the germline at different developmental stages. Homologous chromosomes undergo synapsis during early-mid pachytene.

**(B)** Fluorescent images of germ nuclei in worms expressing ATTF-6::RFP and GFP::Pol II at early-mid pachytene and late pachytene. Each dotted circle marks a germline nucleus. Confocal images (60× objective) are shown as maximum-intensity projections spanning the germline. Arrows indicate the piRNA gene cluster, and arrowheads mark the 5S rDNA-SL1 gene cluster. Scale bar: 2 μm.

**(C)** Fluorescent image of a single germ nucleus at early-mid pachytene. Right, normalized fluorescence intensity profiles of ATTF-6::RFP and GFP::Pol II measured along the dashed line crossing both the large and small ATTF-6::RFP foci. Fluorescence signals were normalized by subtracting background (minimum intensity) and dividing by maximum intensity to yield values from 0 to 100. AU, arbitrary units.

**(D)** Same as (**B**), but with ATTF-6::RFP and GFP::Pol III.

**(E)** Same as (**C**), but with ATTF-6::RFP and GFP::Pol III.

**(F)** Fluorescent images of germ nuclei in worms expressing mCherry::PRDE-1 and GFP::Pol II at early-mid pachytene and late pachytene. Each dotted circle marks a germline nucleus. Confocal images (60× objective) are shown as maximum-intensity projections spanning the germline. Arrows indicate the piRNA gene cluster, and arrowheads mark the small GFP::Pol II foci. Scale bar: 2 μm.

**(G)** Same as (**F**), but with mCherry::PRDE-1 and GFP::Pol III.

**Figure S2. ChIP signals of ATTF-6, RNA Pol II, and RNA Pol III across genome.**

**(A)** Browser view of ATTF-6, Pol II (AMA-1), and Pol III (RPC-1) ChIP-seq signals on chromosome I (Ikegami and Lieb 2013; Wang et al. 2025). ATTF-6 ChIP-seq signals were normalized to the control IP (Wang et al. 2025), and AMA-1 and RPC-1 signals were normalized to the input by subtracting their corresponding control signals (Ikegami and Lieb 2013). The signals represent the average from two biological replicates and are shown in RPKM (Reads Per Kilobase per Million mapped reads).

**(B–E)** Browser views for chromosomes II, III, IV, and X.

**Figure S3. Differential persistence of ATTF-6 and polymerase foci following embryo extrusion.**

**(A, C, E)** Representative time-lapse fluorescence micrographs of extruded embryos expressing GFP::ATTF-6 (**A**), GFP::Pol II (**C**), or GFP::Pol III (**E**). Individual nuclei were identified using H2B::mCherry and are outlined by dashed circles. Regions enclosed by white dashed squares are enlarged in the corresponding insets. Time after extrusion is indicated in minutes. Scale bars, 15 µm; inset scale bars, 5 µm.

**(B, D, F)** Quantification of the average number of GFP::ATTF-6 (**B**), GFP::Pol II (**D**), or GFP::Pol III (**F**) foci per nucleus at 0 and 12 minutes after embryo extrusion. Data were obtained from three independent biological replicates. n represents total number of nuclei analyzed at each replicate.

**Figure S4. Quantification of ATTF6, Pol II, and Pol III foci number in Auxin or RNAi treatment.**

**(A)** Definition of foci patterns used for quantifying the foci in (**B–E**), including two clear foci, multiple small foci, and dissolved foci.

**(B)** Quantification of GFP::Pol II and ATTF-6::RFP foci patterns in embryo nuclei based on the definitions in (**A**). Embryos were dissected from worms treated with or without auxin (1mM IAA). Nuclei from three independent embryos were counted, and the number of nuclei is indicated.

**(C)** Same analysis as in (**B**), but examining GFP::Pol III and ATTF-6::RFP foci.

**(D)** Quantification of GFP::Pol II and ATTF-6::RFP foci patterns in embryo nuclei based on the definitions in (**A**). Embryos were dissected from worms treated with control, *attf-6*, and *snpc-4* RNAi. Nuclei from three independent embryos were counted, and the number of nuclei is indicated.

**(E)** Same analysis as in (**D**), but examining GFP::Pol III and ATTF-6::RFP foci.

**Figure S5. Fraction of ATTF-6::AID::RFP, GFP::Pol II, or GFP::Pol III within the granules over nuclear signals.**

**(A)** Box plot showing the fraction of nuclear fluorescence localized within foci for GFP::Pol II and ATTF-6::AID::RFP without or with 1 mM auxin treatment in the *sun-1p*::TIR1 background. Red dots indicate mean values. Statistical significance was calculated using a two-tailed Student’s *t*-test (**** 0.0001).

**(B)** Same as (A), but showing ATTF-6::AID::RFP and GFP::Pol III. Statistical significance was calculated using a two-tailed Student’s *t*-test (*** 0.001; **** 0.0001).

**(C)** Box plot showing the fraction of nuclear fluorescence localized within foci for GFP::Pol II and ATTF-6::AID::RFP following *control, attf-6, or snpc-4* RNAi treatment. Red dots indicate mean values. Statistical significance was calculated using a two-tailed Student’s *t*-test (*** 0.001; **** 0.0001).

**(D)** Same as (C), but showing ATTF-6::AID::RFP and GFP::Pol III. Statistical significance was calculated using a two-tailed Student’s *t*-test (ns, not significant; ** p ≤ 0.01; **** p ≤ 0.0001).

**Figure S6. Number of ATTF6, Pol II, and Pol III foci per nucleus.**

**(A, C, E)** Violin plots showing the number of foci per nucleus for embryos expressing GFP::ATTF-6 **(A)**, GFP::Pol II **(C)**, or GFP::Pol III **(E)**. Embryos resuspended in M9 buffer were maintained at 20°C or exposed to heat-stress at 32°C. Time (hrs) indicates time post-transfer to a 32°C incubator. Dots represent the median foci number and n represents individual nuclei. Three embryos were quantified per condition. Statistics conducted using the Student’s t-test (ns, not significant; **** p ≤ 0.0001).

(**B**, **D**, **F**) Boxplots showing the fraction of nuclear fluorescence localized within foci for GFP::ATTF-6 (**B**), GFP::Pol II (**D**), or GFP::Pol III (**F**) expressing embryos. Red diamonds indicate the mean. Statistics conducted using the Student’s t-test (ns, not significant; ** p ≤ 0.01; **** p ≤ 0.0001).

## References

Adilardi RS, Dernburg AF. 2022. Robust, versatile DNA FISH probes for chromosome-specific repeats in Caenorhabditis elegans and Pristionchus pacificus. G3 12: jkac121.

Alberti S, Arosio P, Best RB, Boeynaems S, Cai D, Collepardo-Guevara R, Dignon GL, Dimova R, Elbaum-Garfinkle S, Fawzi NL et al. 2025. Current practices in the study of biomolecular condensates: a community comment. Nature communications 16: 7730.

Ashley GE, Duong T, Levenson MT, Martinez MAQ, Johnson LC, Hibshman JD, Saeger HN, Palmisano NJ, Doonan R, Martinez-Mendez R et al. 2021. An expanded auxin-inducible degron toolkit for Caenorhabditis elegans. Genetics 217.

Banani SF, Lee HO, Hyman AA, Rosen MK. 2017. Biomolecular condensates: organizers of cellular biochemistry. Nature reviews Molecular cell biology 18: 285–298.

Batista PJ, Ruby JG, Claycomb JM, Chiang R, Fahlgren N, Kasschau KD, Chaves DA, Gu W, Vasale JJ, Duan S et al. 2008. PRG-1 and 21U-RNAs interact to form the piRNA complex required for fertility in C. elegans. Molecular cell 31: 67–78.

Bird DM, Riddle DL. 1989. Molecular cloning and sequencing of ama-1, the gene encoding the largest subunit of Caenorhabditis elegans RNA polymerase II. Molecular and cellular biology 9: 4119–4130.

Blumenthal T. 1998. Gene clusters and polycistronic transcription in eukaryotes. BioEssays : news and reviews in molecular, cellular and developmental biology 20: 480–487.

-. 2012. Trans-splicing and operons in C. elegans. WormBook: 1–11.

Boehning M, Dugast-Darzacq C, Rankovic M, Hansen AS, Yu T, Marie-Nelly H, McSwiggen DT, Kokic G, Dailey GM, Cramer P et al. 2018. RNA polymerase II clustering through carboxy-terminal domain phase separation. Nature structural & molecular biology 25: 833–840.

Brangwynne CP, Eckmann CR, Courson DS, Rybarska A, Hoege C, Gharakhani J, Julicher F, Hyman AA. 2009. Germline P granules are liquid droplets that localize by controlled dissolution/condensation. Science 324: 1729–1732.

Brenner S. 1974. The genetics of Caenorhabditis elegans. Genetics 77: 71–94.

Brunquell J, Morris S, Lu Y, Cheng F, Westerheide SD. 2016. The genome-wide role of HSF-1 in the regulation of gene expression in Caenorhabditis elegans. BMC Genomics 17: 559.

Cho WK, Spille JH, Hecht M, Lee C, Li C, Grube V, Cisse, II. 2018. Mediator and RNA polymerase II clusters associate in transcription-dependent condensates. Science 361: 412–415.

Cramer P. 2019. Organization and regulation of gene transcription. Nature 573: 45–54.

Das PP, Bagijn MP, Goldstein LD, Woolford JR, Lehrbach NJ, Sapetschnig A, Buhecha HR, Gilchrist MJ, Howe KL, Stark R et al. 2008. Piwi and piRNAs act upstream of an endogenous siRNA pathway to suppress Tc3 transposon mobility in the Caenorhabditis elegans germline. Molecular cell 31: 79–90.

Ding Q, Li R, Ren X, Chan LY, Ho VWS, Xie D, Ye P, Zhao Z. 2022. Genomic architecture of 5S rDNA cluster and its variations within and between species. BMC Genomics 23: 238.

Duronio RJ, Marzluff WF. 2017. Coordinating cell cycle-regulated histone gene expression through assembly and function of the Histone Locus Body. RNA biology 14: 726–738.

Ellis RE, Sulston JE, Coulson AR. 1986. The rDNA of C. elegans: sequence and structure. Nucleic acids research 14: 2345–2364.

Erdel F, Rippe K. 2018. Formation of Chromatin Subcompartments by Phase Separation. Biophys J 114: 2262–2270.

Feng J, Liu T, Qin B, Zhang Y, Liu XS. 2012. Identifying ChIP-seq enrichment using MACS. Nature protocols 7: 1728–1740.

Fire A, Xu S, Montgomery MK, Kostas SA, Driver SE, Mello CC. 1998. Potent and specific genetic interference by double-stranded RNA in Caenorhabditis elegans. Nature 391: 806–811.

Fritsch AW, Diaz-Delgadillo AF, Adame-Arana O, Hoege C, Mittasch M, Kreysing M, Leaver M, Hyman AA, Julicher F, Weber CA. 2021. Local thermodynamics govern formation and dissolution of Caenorhabditis elegans P granule condensates. Proc Natl Acad Sci U S A 118.

Gao Y, Li X, Li P, Lin Y. 2022. A brief guideline for studies of phase-separated biomolecular condensates. Nature chemical biology 18: 1307–1318.

Garrigues JM, Tsu BV, Daugherty MD, Pasquinelli AE. 2019. Diversification of the Caenorhabditis heat shock response by Helitron transposable elements. Elife 8.

Girbig M, Misiaszek AD, Muller CW. 2022. Structural insights into nuclear transcription by eukaryotic DNA-dependent RNA polymerases. Nature reviews Molecular cell biology 23: 603–622.

Grummt I. 2003. Life on a planet of its own: regulation of RNA polymerase I transcription in the nucleolus. Genes & development 17: 1691–1702.

Hnisz D, Shrinivas K, Young RA, Chakraborty AK, Sharp PA. 2017. A Phase Separation Model for Transcriptional Control. Cell 169: 13–23.

Hou X, Zhu C, Xu M, Chen X, Sun C, Nashan B, Guang S, Feng X. 2022. The SNAPc complex mediates starvation-induced trans-splicing in Caenorhabditis elegans. J Genet Genomics 49: 952–964.

Huang SK, Whitney PH, Dutta S, Shvartsman SY, Rushlow CA. 2021. Spatial organization of transcribing loci during early genome activation in Drosophila. Curr Biol 31: 5102–5110 e5105.

Ikegami K, Lieb JD. 2013. Integral nuclear pore proteins bind to Pol III-transcribed genes and are required for Pol III transcript processing in C. elegans. Mol Cell 51: 840–849.

Jackson DA, Hassan AB, Errington RJ, Cook PR. 1993. Visualization of focal sites of transcription within human nuclei. The EMBO journal 12: 1059–1065.

Kamath RS, Fraser AG, Dong Y, Poulin G, Durbin R, Gotta M, Kanapin A, Le Bot N, Moreno S, Sohrmann M et al. 2003. Systematic functional analysis of the Caenorhabditis elegans genome using RNAi. Nature 421: 231–237.

Kasper DM, Wang G, Gardner KE, Johnstone TG, Reinke V. 2014. The C. elegans SNAPc component SNPC-4 coats piRNA domains and is globally required for piRNA abundance. Developmental cell 31: 145–158.

Kechin A, Boyarskikh U, Kel A, Filipenko M. 2017. cutPrimers: a new tool for accurate cutting of primers from reads of targeted next generation sequencing. Journal of Computational Biology 24: 1138–1143.

Kemp JP, Jr., Yang XC, Dominski Z, Marzluff WF, Duronio RJ. 2021. Superresolution light microscopy of the Drosophila histone locus body reveals a core-shell organization associated with expression of replication-dependent histone genes. Mol Biol Cell 32: 942–955.

Kim H, Ishidate T, Ghanta KS, Seth M, Conte D, Jr., Shirayama M, Mello CC. 2014. A co-CRISPR strategy for efficient genome editing in Caenorhabditis elegans. Genetics 197: 1069–1080.

Kipreos ET, van den Heuvel S. 2019. Developmental Control of the Cell Cycle: Insights from Caenorhabditis elegans. Genetics 211: 797–829.

McSwiggen DT, Mir M, Darzacq X, Tjian R. 2019. Evaluating phase separation in live cells: diagnosis, caveats, and functional consequences. Genes & development 33: 1619–1634.

Morton EA, Lamitina T. 2013. Caenorhabditis elegans HSF-1 is an essential nuclear protein that forms stress granule-like structures following heat shock. Aging Cell 12: 112–120.

Muzzopappa F, Hummert J, Anfossi M, Tashev SA, Herten DP, Erdel F. 2022. Detecting and quantifying liquid-liquid phase separation in living cells by model-free calibrated half-bleaching. Nat Commun 13: 7787.

Palacio M, Taatjes DJ. 2022. Merging Established Mechanisms with New Insights: Condensates, Hubs, and the Regulation of RNA Polymerase II Transcription. Journal of molecular biology 434: 167216.

Pastore B, Hertz HL, Price IF, Tang W. 2021. pre-piRNA trimming and 2’-O-methylation protect piRNAs from 3’ tailing and degradation in C. elegans. Cell Rep 36: 109640.

Pastore B, Hertz HL, Tang W. 2022. Comparative analysis of piRNA sequences, targets and functions in nematodes. RNA Biol 19: 1276–1292.

Patel A, Lee HO, Jawerth L, Maharana S, Jahnel M, Hein MY, Stoynov S, Mahamid J, Saha S, Franzmann TM et al. 2015. A Liquid-to-Solid Phase Transition of the ALS Protein FUS Accelerated by Disease Mutation. Cell 162: 1066–1077.

Pazdernik N, Schedl T. 2013. Introduction to germ cell development in Caenorhabditis elegans. Advances in experimental medicine and biology 757: 1–16.

Phillips CM, McDonald KL, Dernburg AF. 2009. Cytological analysis of meiosis in Caenorhabditis elegans. Methods Mol Biol 558: 171–195.

Pombo A, Jackson DA, Hollinshead M, Wang Z, Roeder RG, Cook PR. 1999. Regional specialization in human nuclei: visualization of discrete sites of transcription by RNA polymerase III. The EMBO journal 18: 2241–2253.

Price IF, Hertz HL, Pastore B, Wagner J, Tang W. 2021. Proximity labeling identifies LOTUS domain proteins that promote the formation of perinuclear germ granules in C. elegans. eLife 10.

Price IF, Wagner JA, Pastore B, Hertz HL, Tang W. 2023. C. elegans germ granules sculpt both germline and somatic RNAome. Nat Commun 14: 5965.

Ramírez F, Ryan DP, Grüning B, Bhardwaj V, Kilpert F, Richter AS, Heyne S, Dündar F, Manke T. 2016. deepTools2: a next generation web server for deep-sequencing data analysis. Nucleic acids research 44: W160.

Rippe K. 2022. Liquid-Liquid Phase Separation in Chromatin. Cold Spring Harbor perspectives in biology 14.

Rippe K, Papantonis A. 2025. RNA polymerase II transcription compartments - from factories to condensates. Nature reviews Genetics 26: 775–788.

Robinson JT, Thorvaldsdóttir H, Winckler W, Guttman M, Lander ES, Getz G, Mesirov JP. 2011. Integrative genomics viewer. Nature biotechnology 29: 24–26.

Roeder RG. 2019. 50+ years of eukaryotic transcription: an expanding universe of factors and mechanisms. Nature structural & molecular biology 26: 783–791.

Rogalski TM, Riddle DL. 1988. A Caenorhabditis elegans RNA polymerase II gene, ama-1 IV, and nearby essential genes. Genetics 118: 61–74.

Ruby JG, Jan C, Player C, Axtell MJ, Lee W, Nusbaum C, Ge H, Bartel DP. 2006. Large-scale sequencing reveals 21U-RNAs and additional microRNAs and endogenous siRNAs in C. elegans. Cell 127: 1193–1207.

Stempor P, Ahringer J. 2016. SeqPlots-Interactive software for exploratory data analyses, pattern discovery and visualization in genomics. Wellcome open research 1.

Stortz M, Presman DM, Levi V. 2024. Transcriptional condensates: a blessing or a curse for gene regulation? Commun Biol 7: 187.

Thomas L, Taleb Ismail B, Askjaer P, Seydoux G. 2023. Nucleoporin foci are stress-sensitive condensates dispensable for C. elegans nuclear pore assembly. The EMBO journal 42: e112987.

Vasimuddin M, Misra S, Li H, Aluru S. 2019. Efficient architecture-aware acceleration of BWA-MEM for multicore systems. in 2019 IEEE international parallel and distributed processing symposium (IPDPS), pp. 314–324. IEEE.

Wang YH, Hertz HL, Pastore B, Tang W. 2025. An AT-hook transcription factor promotes transcription of histone, spliced-leader, and piRNA clusters. Nucleic Acids Res 53.

Wansink DG, Schul W, van der Kraan I, van Steensel B, van Driel R, de Jong L. 1993. Fluorescent labeling of nascent RNA reveals transcription by RNA polymerase II in domains scattered throughout the nucleus. J Cell Biol 122: 283–293.

Weick EM, Sarkies P, Silva N, Chen RA, Moss SM, Cording AC, Ahringer J, Martinez-Perez E, Miska EA. 2014. PRDE-1 is a nuclear factor essential for the biogenesis of Ruby motif-dependent piRNAs in C. elegans. Genes & development 28: 783–796.

Weng C, Kosalka J, Berkyurek AC, Stempor P, Feng X, Mao H, Zeng C, Li WJ, Yan YH, Dong MQ et al. 2019. The USTC co-opts an ancient machinery to drive piRNA transcription in C. elegans. Genes & development 33: 90–102.

Yang YF, Zhang X, Ma X, Zhao T, Sun Q, Huan Q, Wu S, Du Z, Qian W. 2017. Trans-splicing enhances translational efficiency in C. elegans. Genome Res 27: 1525–1535.

Zhang L, Ward JD, Cheng Z, Dernburg AF. 2015. The auxin-inducible degradation (AID) system enables versatile conditional protein depletion in C. elegans. Development 142: 4374–4384.

Zhang Y, Liu T, Meyer CA, Eeckhoute J, Johnson DS, Bernstein BE, Nusbaum C, Myers RM, Brown M, Li W. 2008. Model-based analysis of ChIP-Seq (MACS). Genome biology 9: 1–9.

Zheng T, Wake N, Weng SL, Perdikari TM, Murthy AC, Mittal J, Fawzi NL. 2025. Molecular insights into the effect of 1,6-hexanediol on FUS phase separation. The EMBO journal 44: 2725–2740.

