## Supplementary figures and images for "Dynamic RNA Polymerase compartments organize the transcription of gene clusters"

### Supplemental Figures

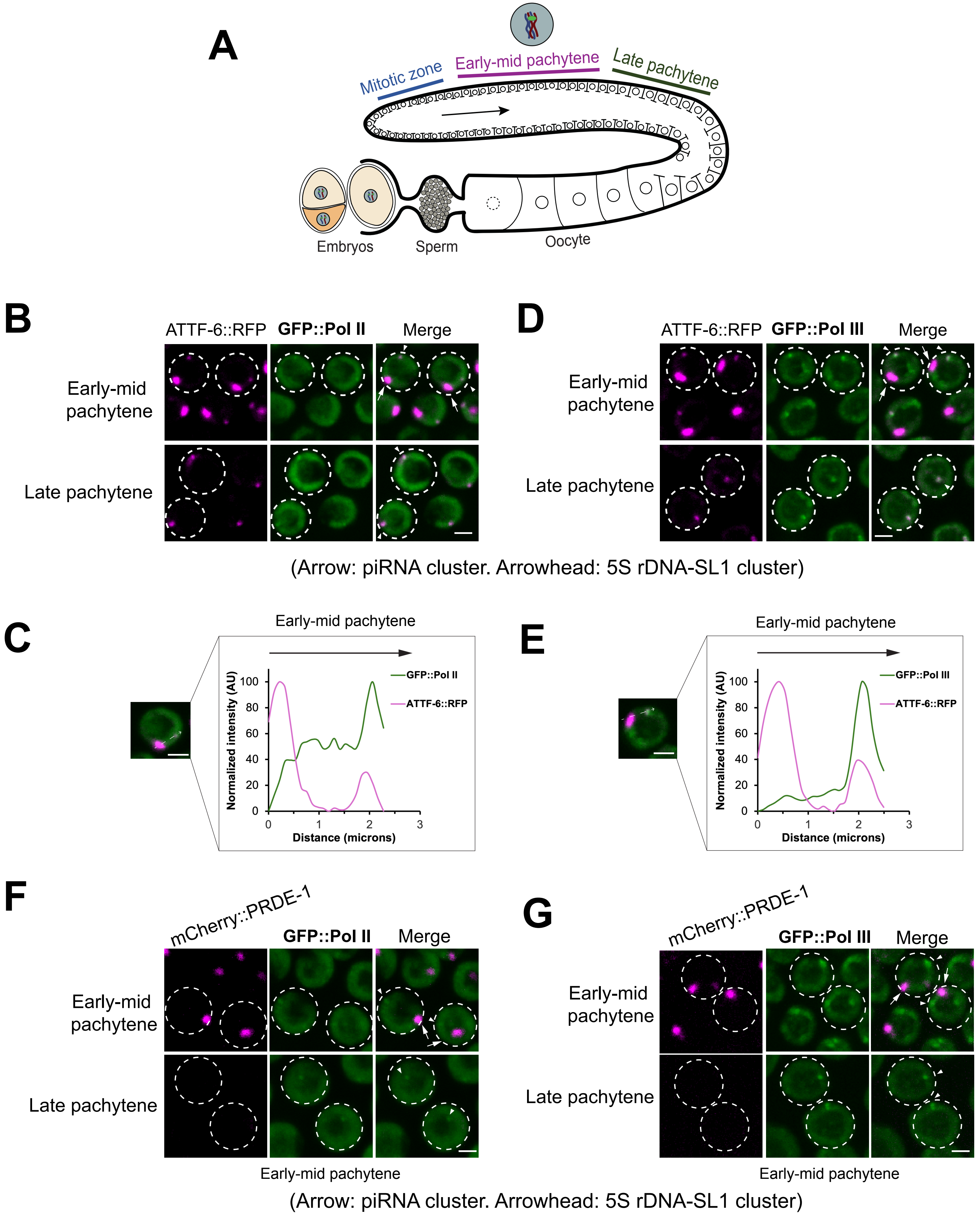

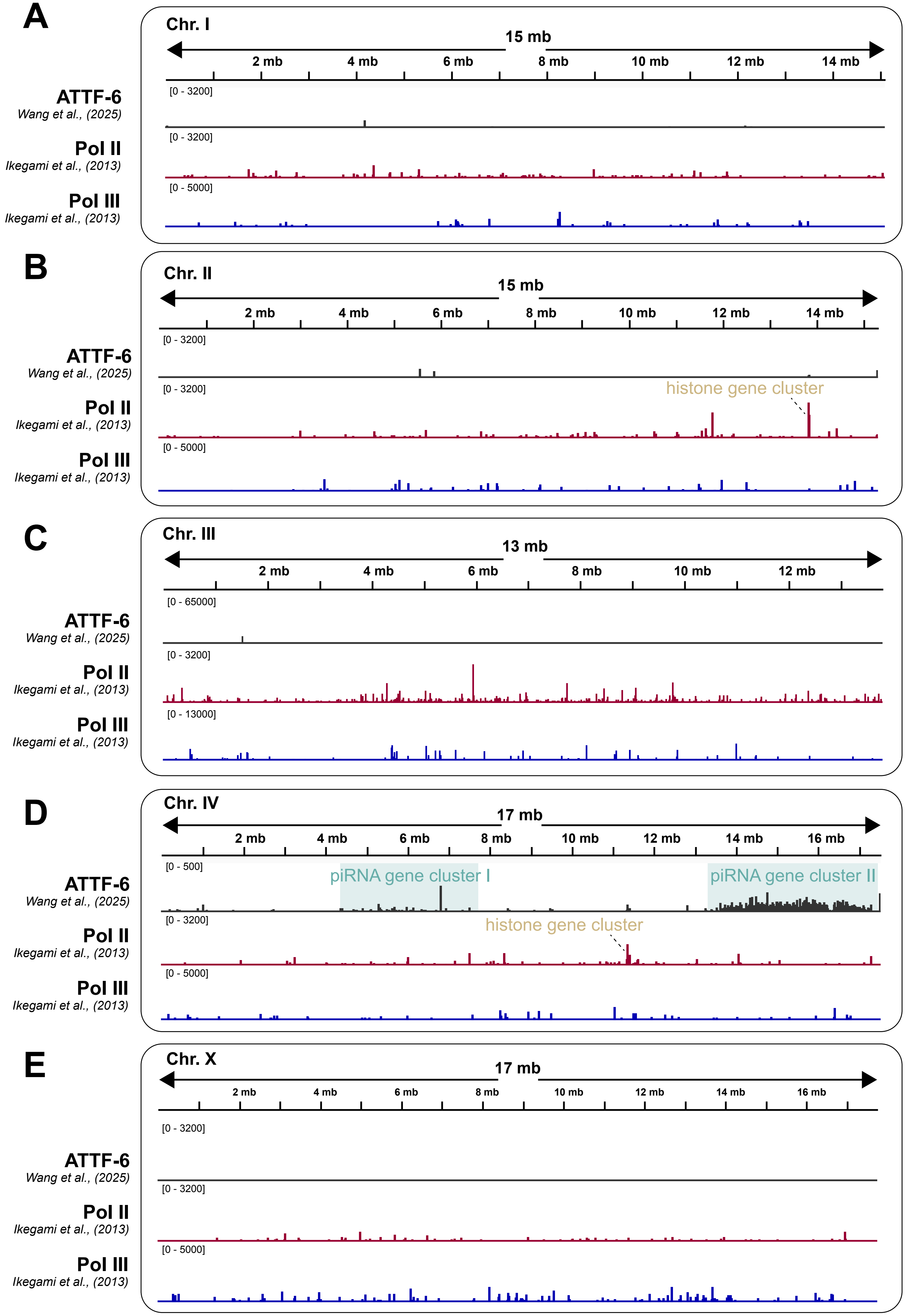

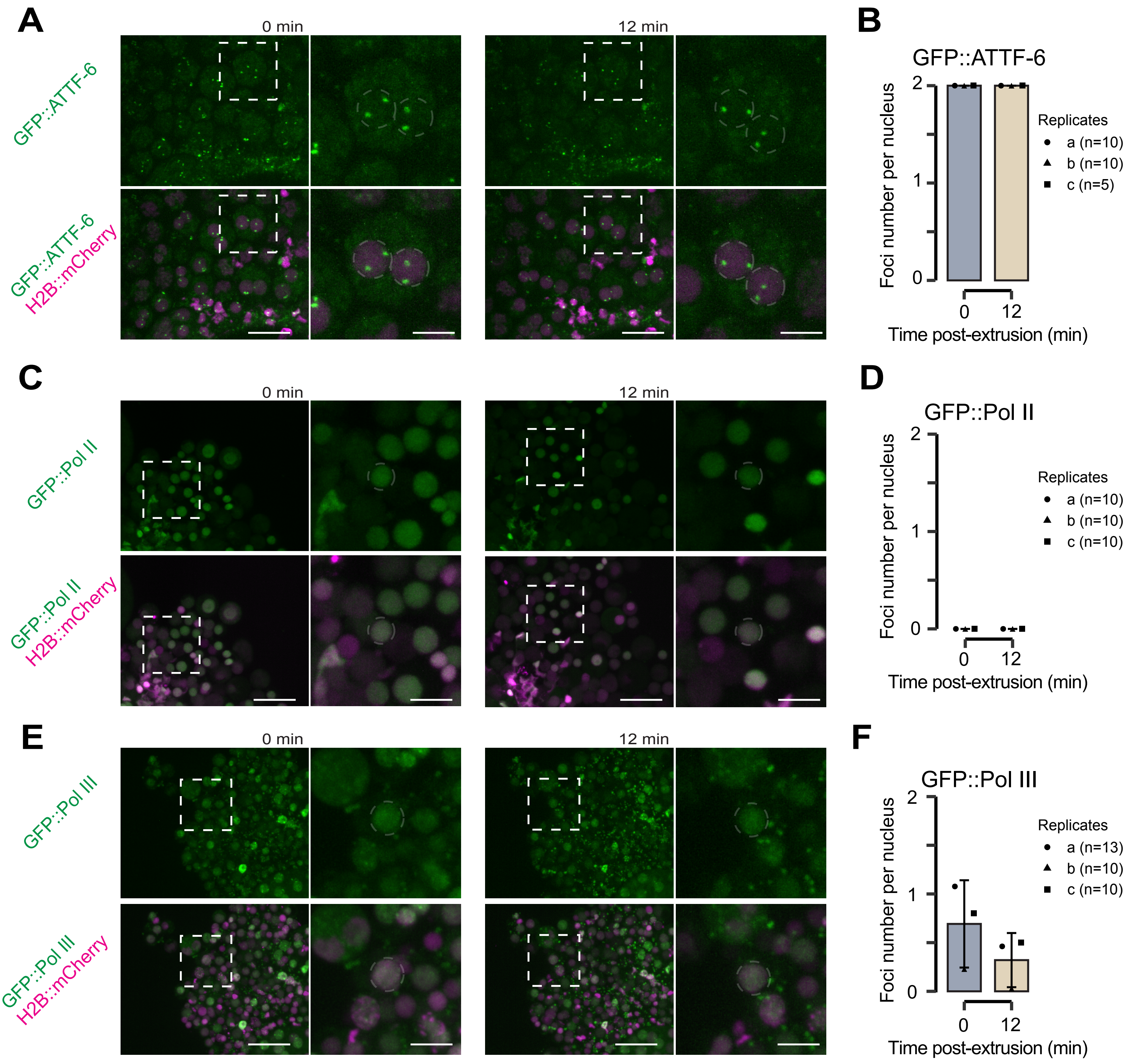

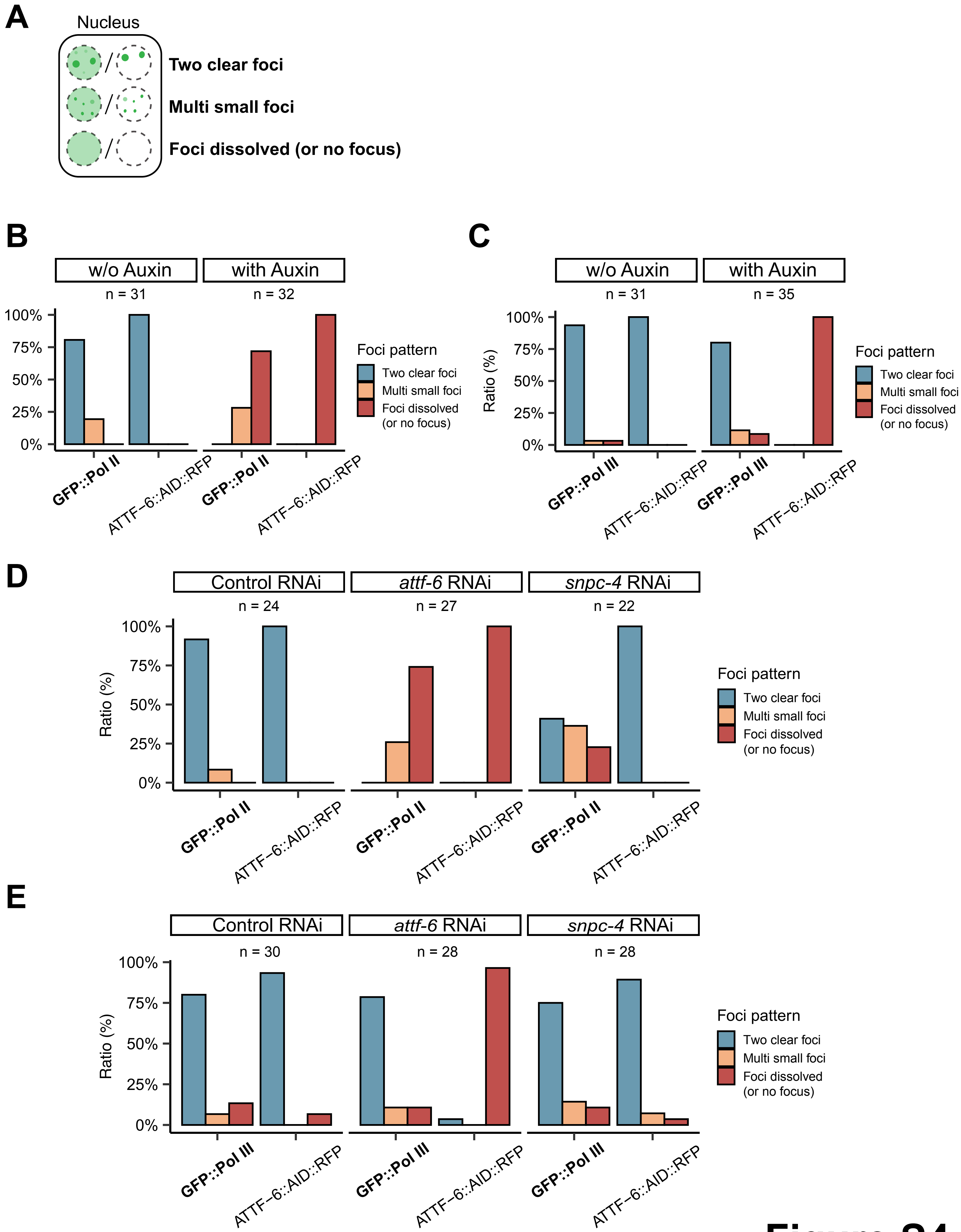

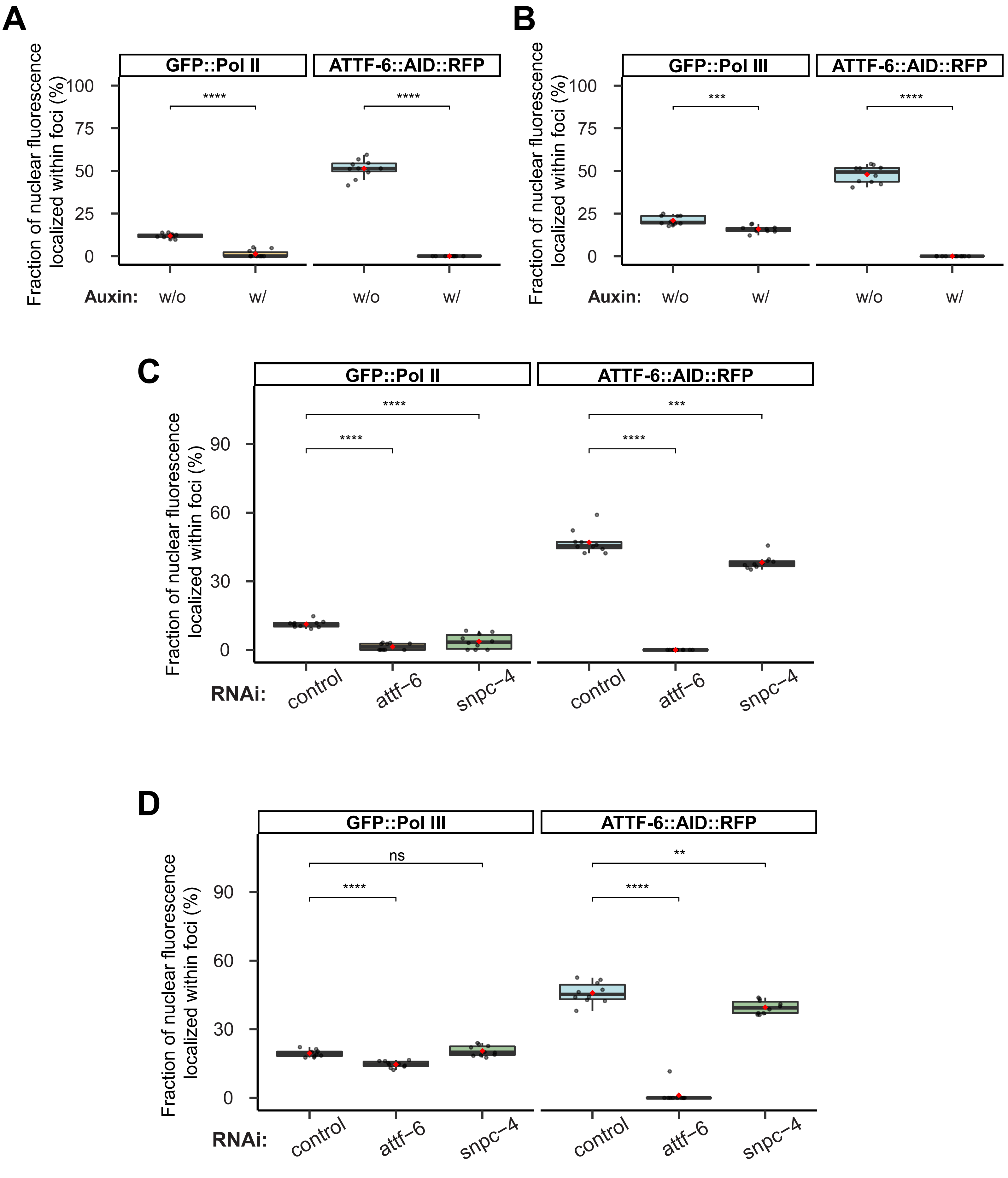

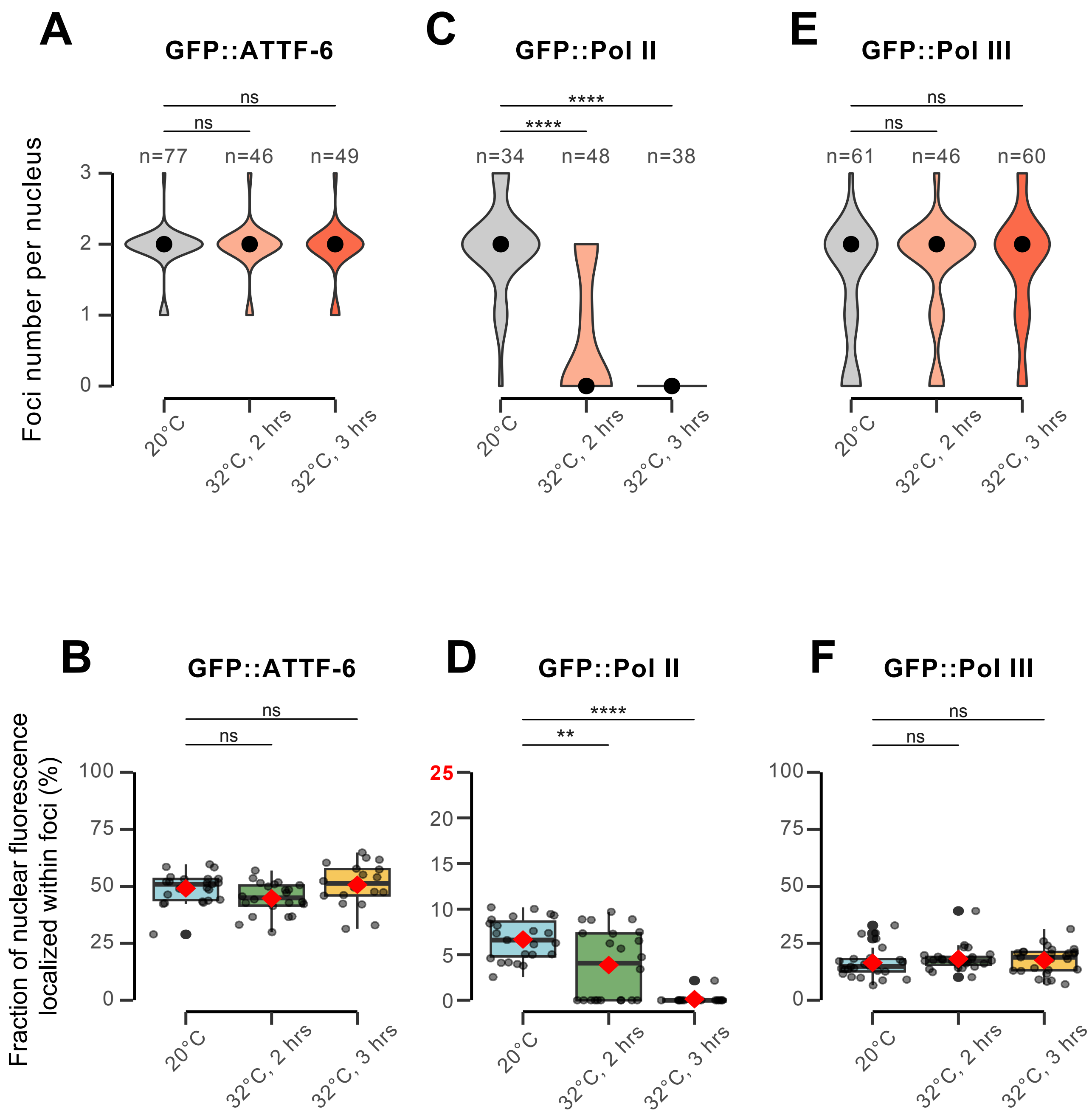
